# Bottom-up fabrication of a multi-component nanopore sensor that unfolds, processes and recognizes single proteins

**DOI:** 10.1101/2020.12.04.411884

**Authors:** Shengli Zhang, Gang Huang, Roderick Versloot, Bart Marlon Herwig, Paulo Cesar Telles de Souza, Siewert-Jan Marrink, Giovanni Maglia

## Abstract

Transmembrane channels and pores have many biotechnological applications, notably in the single-molecule sequencing of DNA. Small synthetic nanopores have been designed using amphipathic peptides, or by assembling computationally designed transmembrane helices. The fabrication of more complex transmembrane devices has yet to be reported. In this work, we fabricated in two steps a multi-protein transmembrane device that addresses some of the main challenges in nanopore protein sequencing. In the first step, artificial nanopores are created from soluble proteins with toroid shapes. This design principle will allow fabricating a variety of nanopores for single-molecule analysis. In the second step one α-subuinit of the 20S proteasome from *Thermoplasma acidophilum* is genetically integrated into the artificial nanopore, and a 28-component nanopore-proteasome is co-assembled in *E. coli* cells. This multi-component molecular machine opens the door to two new approaches in protein sequencing, in which selected substrate proteins are unfolded, fed to into the proteasomal chamber and then identified by the nanopore sensor either as intact or fragmented polypeptides. The ability to integrate molecular devices directly onto a nanopore sensors allows creating next-generation protein sequencing devices, and will shed new lights on the fundamental processes of biological nanomachines.

## Introduction

Membrane-spanning channels and pores have key roles in cellular processes and in biotechnological applications such as nanopore DNA sequencing. Transmembrane proteins have been designed to control the passive transport of molecules across membranes, including a synthetic ion channel^1^, a four-helix divalent metal-ion transporter^2^, a membrane-spanning pore^3^ and a DNA-scaffolded pore^4^. However, the ability to design nanopores with more complex functions has yet to be reported. Such devices would add a new dimension to the protein engineering field, and allow designing next-generation nanopore sensors for biopolymer analysis. Advances in this field, nonetheless, have been hampered by a number of reasons. Usually, molecular machines form multimeric complexes that often require complex post- and co-translational assembly^5^. The latter is particularly challenging because all components must be soluble, unprocessed by proteases, and co-expressed at similar levels. The introduction of artificial transmembrane regions is yet more challenging as it reduces the solubility of the individual component and it can prevent proper assembly. Moreover, the design of the interface between the hydrophobic transmembrane polypeptides and the hydrophilic components has been completely unexplored. Finally, in order to obtain a functional device, the operation of the molecular machine should not occlude the nanopore sensor.

In this work, we fabricated from the bottom-up a transmembrane molecular device that addresses some of the main challenges in nanopore protein sequencing. This 900 kDa, integrated sensor consists of a three-proteins assembled into a 42-component complex, and was made in two steps. In the first step, we designed artificial nanopores from a soluble protein with a toroid shape (**Fig. 1a-d**). The designed synthetic nanopores showed an activity and electrical proprieties identical to natural nanopores. In the second step, the multiprotein 20S proteasome from *Thermoplasma acidophilum*^6^ was incorporated into the artificial nanopore (**Fig. 1e-h**). Two approaches to single-molecule protein sequencing become possible by this design. In the chop-and-drop mode, unfolded proteins are first degraded by the proteasome and the resulting fragment delivered to the nanopore. In the thread-and-read mode, intact substrates are identified as they translocate across the nanopore. Notably, the activity of the proteasome and the unfolding of proteins did not have an influence on the ionic signal.

**Fig. 1.**
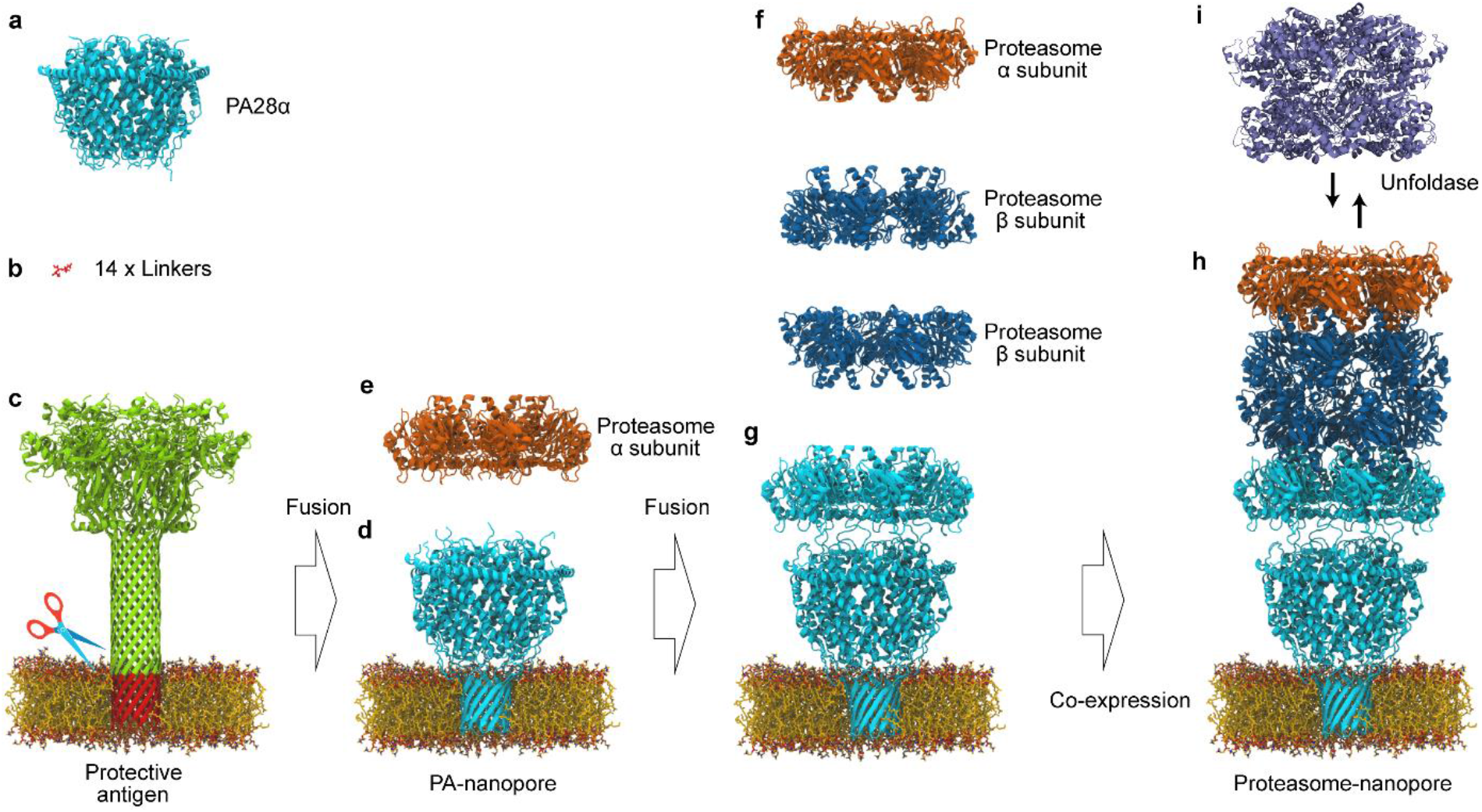
Bottom-up design of a proteasome nanopore. **a**, Structure of mouse PA28α (PDB ID: 5MSJ). **b**, Sticks diagram of the structure of serine-serine-glycine linker. **c**, Ribbon diagram of the structure of anthrax protective antigen (PDB ID: 3J9C). The transmembrane region of the protective antigen is in red. **d**, Structure of the PA-nanopore enhanced by molecular dynamics simulations. PA28α (**a**) was genetically fused to the transmembrane region of the protective antigen (**c**) *via* a short linker (**b**). **e** and **f**, Structure of the *T. acidophilum* proteasome α-and β-subunit (PDB ID: 1YA7). **g**, PA-nanopore was genetically fused to α-subunit of *T. acidophilum* proteasome. **h**, Structure of the designed proteasome-nanopore refined by MD simulations. **i**, Structure of VATΔN (PDB ID: 5G4G).

### Design transmembrane proteins

In cells, PA28α docks onto the 20S proteasome and controls the translocation of protein substrates^7^. Hence, in the first step to build a nanopore-proteasome, we designed a PA28α nanopore. The disorder region of PA28α (from P64 to P100, **Fig. 2a**, red residues) was replaced with the β-barrel transmembrane region (VHGNAEVHASFFDIGGSVSAGF) of anthrax protective antigen^8^. A β-barrel transmembrane region was chosen because it offers high thermodynamic stability. A short flexible linker (SSG) was added to each side of the β-barrel (**Fig. 1a-d, Fig. 2a**, and **Supplementary Fig. 1**) in order to mediate the interaction with the head-groups of the lipid bilayer and to provide a passage for the ion to enter the nanopore.

**Fig. 2.**
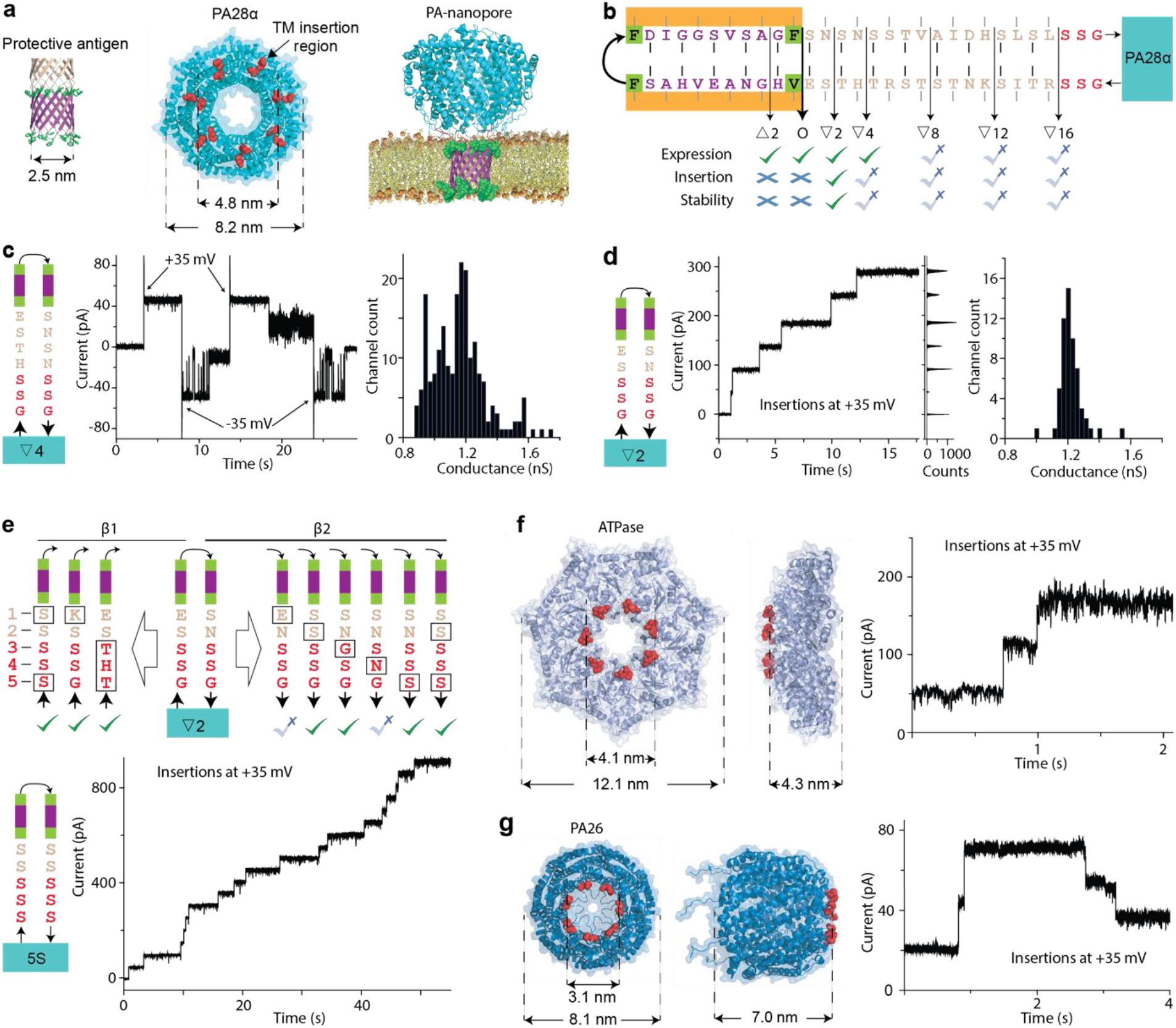
Fabrication and optimization of the artificial nanopores. **a**, Structural representation of the designed nanopores. The β-barrel domain (purple) is introduced within an unstructured loop (red) in PA28α (cyan). The hydrophobic residues anchoring the nanopore to the membrane are indicated in green. The PA-nanopore was generated by molecular dynamics simulations. **b**, Effects of linker length on the nanopore expression in *E. coli*. cells, insertion efficiency and nanopore stability. The side chains that point towards the outside and inside of the barrel are highlighted with grey and black lines, respectively. The first designed nanopore (0) is highlighted with a wider arrow. One deletion mutant (Δ2) and five insertion mutants (∇2, ∇4, ∇8, ∇12, and ∇16) were tested. The sequence of the protective antigen was used as template for the linker. PA28α is shown represented as a cyan rectangle. **c, d** Electrical properties of the functional ∇4 (**c**) and ∇2 (**d**) mutants. On the left is sequence of the mutant, in the middle a typical current trace, and on the right the current histogram corresponding the insertions of multiple pores at +35 mV. **e**, top, linker optimisation tested by substituting several residues with serine. Bottom, Electrical properties of a PA-nanopore with homo-polymeric serine linkers. **f**, ATPase-nanopore formed by introducing the transmembrane barrel elongated with the homo-polymeric serine linkers within a loop (red) at the AAA+ ATPase domain of *Trypanosoma brucei* σ54-RNA polymerase. **g**, Formation of a PA26-nanopore by introducing the β-barrel elongated by the serine linker into a loop in the top face of PA26 from *Aquifex aeolicus*. Electrical data were collected at ±35 mV in 1 M NaCl, 15 mM Tris, pH 7.5, using 10 kHz sampling rate and a 2 kHz low-pass Bessel filter.

Despite the 22 residues of this transmembrane (TM) region are sufficient to span the hydrophobic core of a lipid bilayer, the initial construct did not insert into lipid bilayer (**Fig. 2b**). Since the length of the linker is likely to play an essential role in guiding membrane insertion and in controlling the transmembrane ionic transport, we tested one deletion mutant (Δ2) and five insertion mutants (∇2, ∇4, ∇8, ∇12, and ∇16, **Fig. 2b**). With the exception of Δ2, all variants could insert into the lipid bilayer, although with different efficiency. ∇8, ∇12, and ∇16 showed large current fluctuations, which prevented nanopore characterization (**Supplementary Fig. 2**). ∇4 showed full current blocks and a heterogeneous unitary conductance (**Fig. 2c**). Among all constructs tested, ∇2, which was efficiently expressed and purified (**Supplementary Fig. 3**), produced uniform pores in lipid bilayers with mean unitary conductance (1.17 ± 0.14 nS at -35 mV, 1 M NaCl, 15 mM Tris, pH 7.5, n = 59, **Fig. 2d**). Remarkably, ∇2 (hereafter PA-nanopore) inserted as efficiently and as uniformly as other nanopores found in nature (*e*.*g*. alpha hemolysin^9^), and remained open indefinitely into the lipid bilayer (**Supplementary Fig. 4a**). The individual peptides corresponding to the TM region of anthrax protective antigen could not form nanopores, indicating that a soluble scaffold is required to stabilize the nanopore in lipid bilayers. TEM images showed that the PA-nanopore assemble into oligomers (**Supplementary Fig. 5a**).

In order to validate this design principle, we first generalised the linker sequence. We found that amino acid substitutions with serine residues were well tolerated (**Fig. 2e**). Hence, the β-barrel sequence elongated by linkers containing five serine residues was introduced in two additional soluble proteins. The first was a AAA+ ATPase domain of *Aquifex aeolicus*, which activates the transcription σ54-RNA polymerase^10^ (**Fig. 2f** and **Supplementary Fig. 1**). The second was PA26, a proteasome activator from *Trypanosoma brucei*^11^ (**Fig. 2g** and **Supplementary Fig. 1**). Despite both proteins had a different diameters and surface residues, they inserted into lipid bilayers forming open nanopores, indicating that the β-barrel transmembrane domain and a five-amino acid hydrophilic linker allows a generic method to introduce soluble proteins into lipid bilayers for biopolymer analysis. Pa28α was preferred to PA26 as the latter occasionally closed in planar lipid bilayers (**Fig. 2g**).

### Functional properties of the optimized artificial pore

Molecular dynamics (MD) simulations were performed on the PA-nanopore to better understand the electrostatic and hydrophobic Interactions between the nanopore and the lipid bilayer. As shown in **Fig. 2a** and **Fig. 3a**, two rings of hydrophobic residues anchor the TM region to the hydrophobic edges of the bilayer, while alternated residues with aliphatic side-chains interface the core of the bilayer. The lumen of the pore is kept hydrated by hydrophilic residues. As expected, the hydrophilic side-chain of the linker residues are interacting with the charged head groups of membrane lipids. Inside the β-barrel region, the PA-nanopore showed an increased occupancy of cations compared to anions (**Supplementary Fig. 6**). Similar to other β-barrel nanopores such as αHL^9^, PA-nanopores showed an asymmetric current–voltage (*I*–*V*) relationship (**Fig. 3b**,**c**). Ion-selectivity measurements using asymmetric NaCl concentrations (0.5 M/*cis* and 2 M/*trans*) confirmed the nanopore is cation selective (P_Na_^+^/P_Cl_^-^ = 1.76 ± 0.20, **Fig. 3d**). Here and throughout the manuscript, uncertainties indicate the standard deviations obtained from at least three experiments. The correct folding of the PA-nanopore in the lipid bilayer was characterized using cyclodextrins (CDs), circular molecules that binds to β-barrel nanopores^12^. α-CD, β-CD and γ-CD were added to the *cis* side of the artificial nanopore and the magnitude of the ionic current associated with a blockade (*I*_B_) was measured. It was reported that only β-CD and γ-CD can block anthrax protective antigen nanopores,^13^presumably because α-CD translocates too quickly to be observed. Accordingly, no current blockades were observed when α-CD was tested,. By contrast, β-CD and γ-CD showed characteristic blockades (**Fig. 3e** and **Fig. 3f**). Finally, the ability of the nanopore to identify peptides was tested using angiotensin I (10 amino acids, net charge 0) and dynorphin A (17 amino acids, net charge +4). We found that the two peptides induced blockades which could be easily distinguished using several parameters, including the residual current and the duration of the current blockades (**Fig. 3g** and **Fig. 3h**). Peptides smaller than angiotensin II (8 amino acids, net charge 0) could not be observed by nanopore recordings (**Supplementary Fig. 7**), thus providing the detection limit of oligopeptide detection using PA-nanopore.

**Fig. 3.**
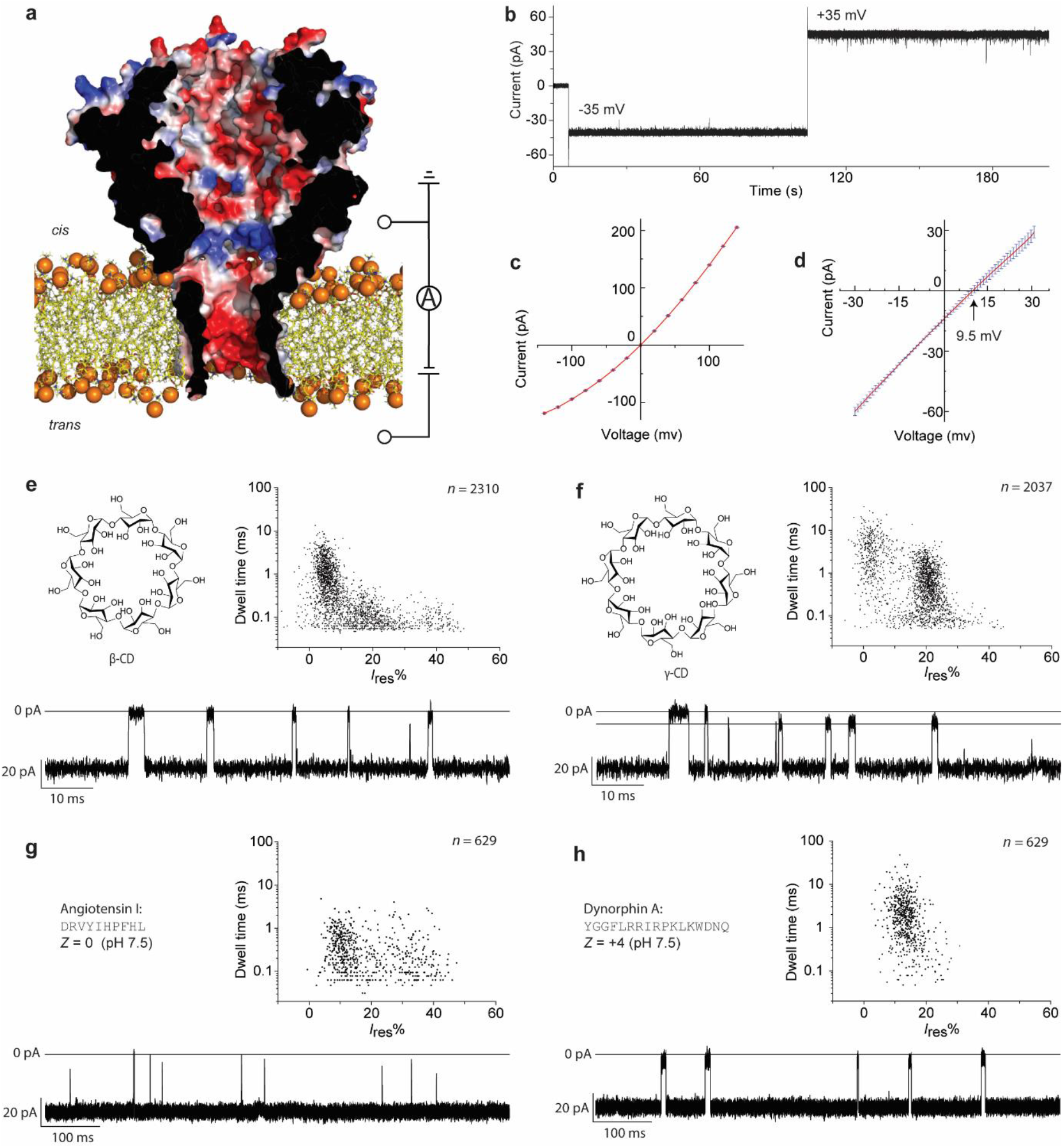
Electrical properties of PA-nanopre and discrimination of substrates. **a**, Cut-through of a surface representation of PA-nanopore. The pore is coloured according to the vacuum electrostatic potential as calculated by PyMOL on the final snapshot of the multiscale MD model. **b**, A typical current trace recorded of a single PA-nanopore at ±35 mV. **c**, Current– voltage (*I*–*V*) characteristics of three different nanopores. **d**, Reversal potential measured using asymmetric ion concentrations (*trans*:*cis*, 2.0 M NaCl: 0.5 M NaCl), showing that the pore is cation-selective, as expected from the electrostatic potentials of the nanopore lumen. The PA-nanopore was added to the *trans* side. **e**, Chemical structure of β-CD (left), scatter plots of *I*_res_% versus dwell time (right), and representative trace of β-CD blockades (below). **f**, Chemical structure of γ-CD (left), scatter plots of *I*_res_% versus dwell time (right), and representative trace of γ-CD (below). **g**, Peptide sequences and net charge of angiotensin I (left), scatter plots of *I*_res_% versus dwell time (right), and representative trace (below). **h**, Peptide sequences of dynorphin A (left), scatter plots of *I*_res_% versus dwell time (right), and representative trace (below). The PA-nanopore and analytes were added to the *cis* side. Electrical data were collected at ±35 mV in 1 M NaCl, 15 mM Tris, pH 7.5, using 10 kHz sampling rate and a 2 kHz low-pass Bessel filter.

### Building a proteasome-nanopore

In the second and final step, the PA-nanopore is fused with the 20S proteasome from *Thermoplasma acidophilum*. The latter is made by four stacked rings composed of 14 α- and 14 β-subunits (**Fig. 1e** and **Fig. 1f**)^6^. The two-flanking outer α-rings allow for the association of the 20S proteasome with several regulatory complexes^14^, among which is proteasome activator PA28α (**Fig. 1a**)^15^. We found, however, that when the proteasome was added to the *cis* side of individual PA-nanopore, no clear interaction was observed. This is most likely because the high ionic strength used (1 M NaCl) does not allow such interaction^16^. The crystal structure of the *Thermoplasma acidophilum* proteasome in complex with PA26 from *Trypanosoma brucei*^11^, a homolog of PA28α, shows that the carboxy-terminal tails of PA26 slide into a pocket on the 20S proteasome, near the amino-terminus of the α-subunit (**Fig. 4a**). Hence, we fused the C-terminal of PA28α (Y249 in PA28α corresponding to S231 in PA26, **Fig. 4a**) with L21 of a proteasome α-subunit, in which the first 20 residues are removed (Δ20-α-subunit), leaving the proteasome gate open towards the PA-nanopore. The formation of the proteasome requires co-assembly of the α and β-subunits. Thus, PA-nanopore fused to proteasome Δ20-α-subunit (PA28-αΔ20 nanopore) containing a C-terminal His-tag, a second opened proteasomal subunit (αΔ12, where the first 12 residues are removed allowing the fast degradation of unfolded substrates without the need for a proteasome activator^17^) containing a C-terminal Strep-tag, and the proteasome β-subunit were co-expressed in *E. coli* cells using a two-vector system (**Extended Methods in Supporting Information** and **Fig. 4b**). Although all proteins could be expressed, SDS analysis indicated that the proteasome-nanopore was proteolytically cleaved inside *E. coli* cells during expression (**Supplementary Fig. 8**). Proteolysis was prevented by optimizing the linker length between the PA-nanopore and α-subunit and by introducing a poly-histidine at the N-terminus of the PA-pore. The latter was introduced to shield the linker between subunits from cellular proteases (**Supplementary Fig. 8**). A co-assembled proteasome-nanopore (mutant 8, **Supplementary Fig. 8)** was then purified in two steps by affinity chromatography (**Extended Methods in Supporting Information** and **Fig. 4b**). SDS-PAGE, native PAGE, and TEM confirmed the successful assembly of the multi-protein complex (**Fig. 4d and Supplementary Fig. 5d**). Activity assays revealed that the proteasome-nanopore was active, with the proteolytic activity increasing with the temperature, and decreasing with the salt concentration (**Supplementary Fig. 9**). The transmembrane proteasome inserted efficiently in lipid bilayers and spontaneous release from the lipid bilayer was not observed (**Supplementary Fig. 4b**). The proteasome-nanopores showed low-noise current recordings at negative applied potentials (**Fig. 4c**). However, at applied potentials higher than +20 mV the proteasome-nanopore often gated (**Supplementary Fig. 10**). This electrical behaviour allowed distinguishing the proteasome-nanopore from PA-nanopore. The *I-V* curve of the proteasome-nanopore was similar to that of PA-nanopore (**Supplementary Fig. 11**), suggesting that the transmembrane region was unchanged and that the proteasome above the nanopore does not influence the ionic signal (see additional discussion is supporting information).

**Fig. 4.**
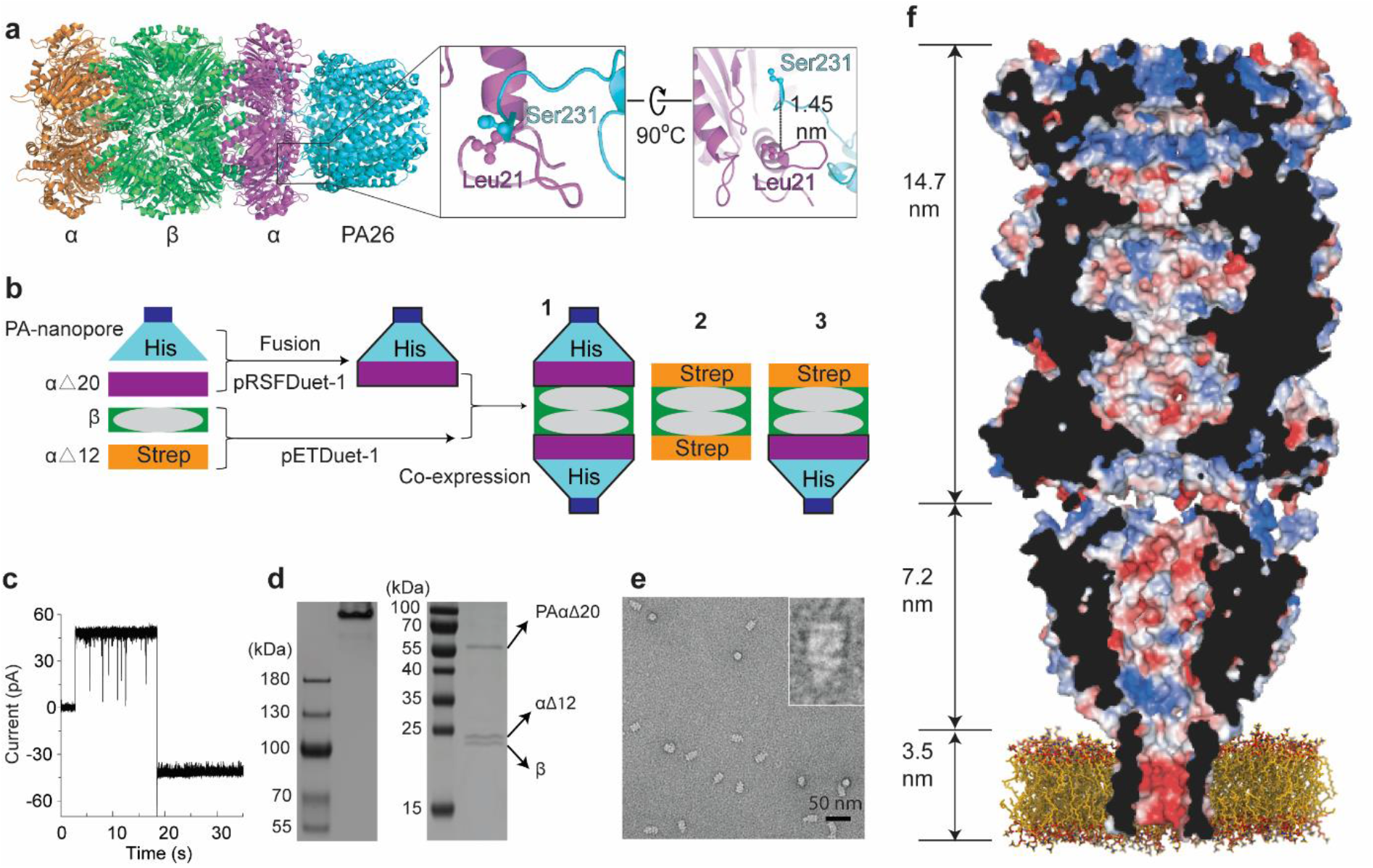
Design of the artificial proteasome-nanopore. **a**, Structure of the *T. acidophilum* proteasome in complex with PA26. PA26 is coloured cyan, the proteasome α-subunit is colored orange and magenta, and the β-subunit is coloured green. The C-terminal of PA26 (S231) is near L21 of the α-subunit. **b**, A representation of the reconstitution of the artificial proteasomal nanopore. To obtain complex **3**, two separate vectors were used to express the four proteins. The PA-nanopore was fused to the proteasome α-subunit and contained a His-tag. The protein was co-expressed with the untagged β-subunits and a second α-subunit containing a Strep-tag. His-tag affinity chromatography was used to co-purify complex **1** and **3**. Then a Strep-Tag affinity chromatography was used to purify **3. c**, Behaviour of a single pore at ±35 mV in 1 M NaCl, 15 mM Tris, pH 7.5, using 10 kHz sampling rate and a 2 kHz low-pass Bessel filter. **d**, Native PAGE (left, ∼5 µg) and SDS-PAGE (right, ∼2 µg) analyses of the purified complex **3**. SDS-PAGE revealed the presence of three unique bands corresponding well the molecular weights of PAαΔ20, proteasome αΔ12-subunit, and proteasome β-subunit (52.7, 25.8, and 22.3 kDa). The native PAGE showed that PAαΔ20, proteasome αΔ12-subunit, and proteasome β-subunit a stable complex **3. e**, TEM image of the proteasome-nanopore. **f**, Cut-through of a surface representation of proteasome-nanopore enhanced by molecular dynamics simulations and coloured (blue, positive; red, negative) according to the vacuum electrostatic potential as calculated by PyMOL.

### Real-time protein processing

The activity of the transmembrane proteasome was tested using substrates containing a C-terminal ssrA tag, which mediates the interaction with VAT (Valosin-containing protein-like ATPase of *Thermoplasma acidophilum*)^18^. The latter is a processive hand-over-hand unfoldase that pulls on two extended residues of substrates through the proteasome chamber^19^. Here, we used VATΔN^20^, in which the first 183 amino acids of the N-terminal domain corresponding to a regulatory domain were deleted. VATΔN displays higher unfolding activity than wild type VAT^20^. We tested two substrates. The first, named S1 (123 amino acid residues), was designed to be unstructured. It contained an ssrA tag followed by four stretches of 15 serine residues, each flanked by 10 arginine residues and three hydrophobic residues (FYW, **Supplementary Fig. 12**). The polyarginine residues were introduced to induce the electrophoretic transport across the nanopore, while the hydrophobic residues are ideal targets for the proteolytic activity of the proteasome. The ssrA tag was introduced to allow recognition with VATΔN. ‘Superfolder’ green fluorescent protein (GFP)^21^, was modified by adding 10 arginines and a ssrA tag at the C-terminus (**Supplementary Fig. 12**) was also tested. GFP was selected because of it high stability towards temperature denaturation (melting temperature 78 °C^22^) and chemotropic agents (unfolding at more than 4M Gu.HCl^23^), hence providing a good model system to test the limit of the proteasome-nanopore to unfold proteins. The activity of VATΔN was optimized using bulk assays (**Supplementary Fig. 13**). Hereafter, experiments performed at 40 °C in 1 M NaCl, 15 mM Tris-HCl, pH 7.5, 20 mM MgCl_2_ solutions. Tests were performed using either proteolytic active or inactive proteasome-nanopores. In the latter, the amino-terminal threonine 1 in the active site was replaced with alanine^5,24^.

### Thread-and-read

The nanopore-proteasome allows two approaches to protein sequencing and identification. In the first, named thread-and-read, proteins are unfolded by VAT and thread across inactivated proteasome. The linearised polypeptide then translocate across the nanopore by the action of the electroosmotic flow. We tested this approach using S1 and GFP. The addition of folded GFP induced no blockades. By contrast, the addition of unfolded S1 induced many blockades that showed a residual current (*I*_res_%, defined as the percent ratio between the blocked nanopore current the open nanopore current) of 7.3 ± 0.1%, which reflect the translocation of unfolded S1 through the nanopore. The blockades were either short (dwell time: 0.30 ± 0.01 ms,) or second-long (**Supplementary Fig. 14**), the latter most likely indicating the occlusion of the proteasome chamber by the substrate. When an equal concentration of VATΔN and ATP (2.0 mM) was added to the solution, S1 blockades became longer (6.64 ± 0.21 ms) and the *I*_res_% increased about ten-fold (70.2 ± 1.0, **Fig. 5a, Supplementary Fig. 15**-**16**), reflecting the VAT-assisted and stretched translocation of S1 across the nanopore. In the presence of VATΔN and ATP, GFP blockades were also observed (**Fig. 5a**), indicating that the unfoldase linearised and fed the substrate protein through the nanopore. GFP blockades were not observed in the absence of either VATΔN or ATP, or when PAN^25^ unfoldase or WT-VAT (WT-VAT contains an N-terminal domain that inhibits its unfolding activity^20^) were used instead of VATΔN (**Supplementary Fig. 17**). When the ATP concentration was increased to 6.0 mM, the average dwell time of GFP blockades decreased to 2.4 ± 1.7 ms (**Supplementary Fig. 18-20**), indicating that VATΔN is capable of feeding the polypeptide through the nanopore at a speed that can be tuned by the concentration of ATP. Contrary to S1 blockades, VAT-assisted GFP blockades showed *I*_res_% close to zero, suggesting the refolding of the substrate protein inside the proteasomal chamber before translocating across the nanopore. In the presence of 1 M urea, GFP current blockades became similar to the blockades induced by unstructured S1 (dwell time: 7.8 ± 1.7 ms, *I*_res_%: 70.3 ± 0.9, **Fig. 5a, Supplementary Fig. 16** and **Supplementary Fig. 21**), indicating that urea most likely prevented the partial refolding of the substrate allowing a stretched translocation across the nanopore. Importantly, the unfolding of the protein above the nanopore did not affect the ionic signal (supporting information for a more detailed discussion), indicating that the signal only reflects the transport of the polypeptide across the nanopore.

**Fig. 5.**
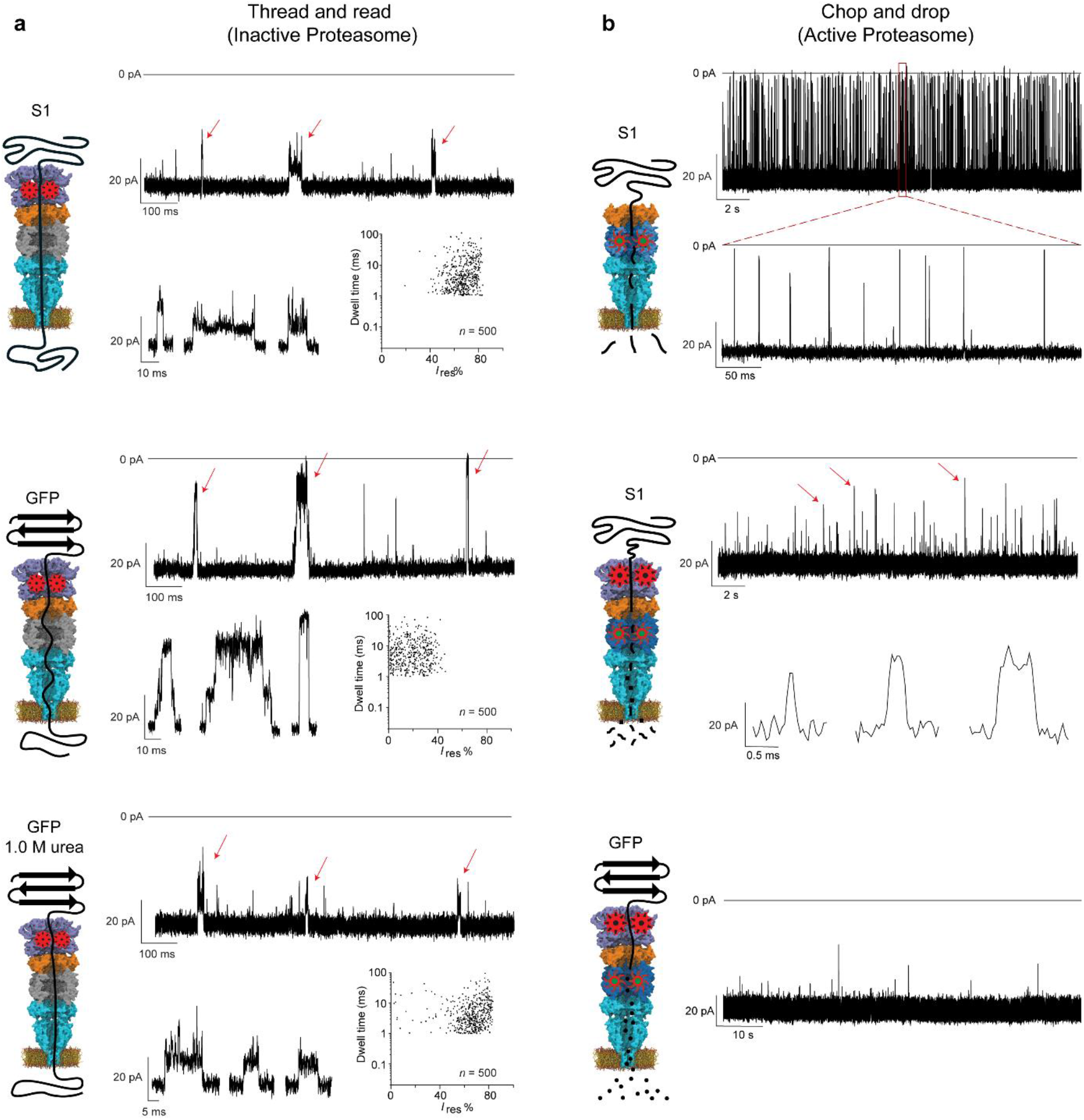
Controlled translocation through the proteasome-nanopore. a, Thread -and-read. Typical current trace and scatter plots of the average *I*_res_% versus dwell time provoked by the translocation of polypeptides through an inactive proteasome-nanopore mediated by VATΔN in the presence of 2.0 mM ATP. From top to bottom: S1 (20.0 µM and 20.0 µM VAT, 23 independent nanopore experiments), GFP (5.0 µM and 5.0 µM VATΔN, 42 independent experiments), GFP in 1 M urea (5.0 µM and 5.0 µM VATΔN, 13 independent experiments). The proteasome-nanopore and substrates were added to the *cis* side. Data were collected at 40 °C and -30 mV in 1 M NaCl, 15 mM Tris, pH 7.5, using a 10 kHz low-pass Bessel filter with a 50 kHz sampling rate. The traces were then filtered digitally with a Gaussian low-pass filter with a 5 kHz cut-off. **b, Cut-and-drop**. Typical current traces provoked by the transport of oligopeptides across an activated proteasome. From top to bottom. Unassisted S1 translocation (>50 independent experiments). VATΔN and ATP assisted S1 transport induce fragmentation into small peptides, which fast transport is seldomly observed (23 independent experiments). VAT assisted GFP-ssrA proteolytic cleavages produce peptides that are too short to be detected by the nanopore (15 independent experiments). Data were collected at 40 °C and -30 mV in 1 M NaCl, 15 mM Tris, pH 7.5, using a 10 kHz low-pass Bessel filter with a 50 kHz sampling rate. The traces were then filtered digitally with a Gaussian low-pass filter with a 5 kHz cut-off.

In a previously published work, target proteins elongated by a C-terminal polypeptide were partially threaded across a nanopore and then forcefully translocated by a ClpX unfoldase added on the opposite (trans) side of the nanopore^26,27^. Crucially, however, the complex current signature arised from the unfolding process at the mouth of the nanopore and not from the transport of the polypeptide across the nanopore, hence fundamentally preventing the recognition of individual amino acids. Further, only folded proteins could be analysed using this method; which is a limitation, considering that unfolded or partially unfolded proteins constitute 40% of Eukaryotes proteins^28,29^. By contrast, in the cis-threading described here, the ionic signal does not reflect the unfolding of proteins but only the passage of the unhindered polypeptide across the nanopore. Therefore, providing a recognition site for individual amino acids can be introduced within the nanopore, this molecular system is compatible with the *de novo* sequencing of proteins.

### Chop-and-drop

The enclosed architecture of the VATΔN-proteasome-nanopore allows a fundamentally new approach in nanopore protein sequencing, in which the proteasome cleaves an unfolded protein and the resulting peptides are recognized as they translocate the nanopore. We recently showed that the ionic signal from peptide blockades to a FraC nanopore relate directly to the volume of the peptide^30,31^. Hence, a proteasome-nanopore might be used as the equivalent of a single-molecule mass spectrometer. We tested this approach using S1 and GFP. When an active proteasome-nanopore was used, the second-long blockades observed during unassisted S1 translocation disappeared, while the short blockades became faster (0.20 ± 0.01 ms, **Fig. 5b** and **Supplementary Fig. 22a**). These results suggest, therefore, that the proteasome processes the substrates as they translocate across the nanopore. When VATΔN and ATP were added in solution, more spaced and shorter blockades were observed (**Fig. 5b** and **Supplementary Fig. 22b**), indicating that the reduced speed of polypeptide threading across the proteasomal chamber allowed the degradation of S1 into smaller peptides that are quickly transported across the nanopore. Accordingly, when GFP was tested under the same conditions yet fewer blockades were observed. The size-limit of peptide detection of the PA-nanopore is ∼8 amino acids (**Supplementary Fig. 6**), while the proteasome has been shown to produce peptide fragments between ∼6 and ∼10 amino acids depending on the protein substrate^32^. In turn this suggests that the slower unfolding of GFP compared to the unstructured S1 allowed for a more efficient proteolysis of the substrate into yet smaller peptides (**Fig. 5b**), which are transported across the nanopore too quickly to be observed.

In conclusion, this work describes a strategy to build nanopores with advanced functionalities, which was used to fabricate from the bottom up a 42-protein component nanopore sensor. This multi-protein molecular machine opens the door to two new approaches in protein sequencing. In the thread-and-read approach, the proteasome is inactivated and polypeptide chains pass intact across the nanopore. Importantly, the current signal reflected the polypeptide transport across the nanopore, but not the unfolding of the protein. Therefore, if the nanopore size can be reduced to sub-nanometer dimension, for example by introducing an alpha-helical transmembrane domain such as that of heptameric FraC^30^, single amino acid recognition should become possible^33^. In the cut-and-read mode selected proteins are digested into smaller peptides within the proteasomal chamber and individual peptides are recognized in sequence by specific current blockades. At present the β-barrel nanopore is too large for proteasomal peptide characterisation. However, we have previously shown that FraC nanopores can identify peptides composed by just three amino acids^30^. Hence, if either the size of peptide cleavage ^24,34^ or the proteolytic pattern of the proteasome can be tuned^35^, this approach should become suitable for real-time and single-molecule protein identification.

## Aknowlegements

This work is financially supported by ERC consolidator grant (number: 726151).

## Author contribution

S.Z. and G.M. designed the experiments. G.M. supervised the project. S.Z. performed the experiments and data analysis. B.B., P.S., and S.M. conducted the simulation work. G.M. and S.Z. wrote the paper. All authors discussed the results, and commented on the manuscript.

## Supporting information

### Additional discussion

#### VATΔN activity does not influence ionic transport across the PA-nanopore

We found that the introducing the proteasome above the PA-nanopore and the motions of VATΔN attached to the proteasome did not have an effect on the ionic signal across the nanopore (**Fig. 4c**, main text). Indeed, the *I-V* curve and the electrical noise of the proteasome-nanopore was almost identical to that of PA-nanopore (**Supplementary Fig. 11**). This finding might not be entirely surprising, as proteins moving polymers on the *cis* side of the nanopore usually do not have an effect on the ionic signal. This has been the case for the helicase-assisted or polymerase-assisted motion of DNA across a range of nanopores, including αHL^9^, MspA^36^ and CsgG^37^. In addition, previous work with αHL nanopores suggests that the *cis* entry does not have a strong influence on the ionic current passing through the nanopore. The mutation K8A, which is right at the *cis* entry of the nanopore does not change the *I-V* curve^9,38^, while the same substitution on the β-barrel has a strong effect on the conductance^9^. In addition, the attachment of DNA aptamer at the *cis* entry of an αHL nanopore does not change its conductance significantly^39^. Furthermore, when thrombin was bound to the aptamer, the ionic current increased rather than decreased, and only by 0.5 pA^39^. Multiple attachment possibilities and different aptamer: proteins conjugates were tested and none gave a signal (Rotem, personal communication). Hence, it is likely that ions enter the β-barrel of the αHL nanopore by alternative pathways. Interestingly, αHL has small side entrances open to the solvent, which might allow ions from the solution to enter the β-barrel. A previous work found that the noise in the conductance of the nanopore is dependent on the pH of the solution with an effective pKa of 5.5^40^. This value corresponded well to the pKa of histidine residues, which are near the small side entrances of αHL.

As for αHL, ions can be transported through the transmembrane region of PA-pore by alternative paths, as shown in the model of the PA-pore (**Supplementary Fig. 22**). Most notably, MD simulations revealed that the connection between the α-subuinit and the PA-pore is particularly open to the solution. An additional path might be the connection between the β-strand and the PA28α and throughout the PA28α structure, or through other part of the nanopore (**Supplementary Fig. 22**). Additional evidences suggest that binding of VATΔN to the proteasome does not have an effect on the electrical signal. During VATΔN-assisted translocation of substrates, the baselines of the proteolytically-active proteasome-nanopore and that of the proteolytically-inactive proteasome-nanopore were identical. Note that during GFP proteolysis spikes from the fast translocation of peptides across the nanopore were observed (**Fig. 5**), while during the transport of intact polypeptides long events were observed (**Supplementary Fig. 15-20**). Since in both mode of action VATΔN actively threads protein across the proteasomal chamber, these results indicate that the action of VATΔN does not influence the ionic signal. In addition, the baseline did not change during other control experiments in which substrate translocation events were not observed, further confirming that the VATΔN activity does not influence the ionic signal. Finally, In the absence of urea, the signal from VATΔN-assisted translocation of S1 and GFP were different, while in the presence of 1M urea the signal from both proteins becomes alike. All these evidences suggest that the link region of the PA-nanopore most likely allow the entry of ions from solution, thus minimizing the effect of the proteasome and the unfoldase at the entry of the nanopore.

## Methods

### General materials

Oligonucleotides and gBlock gene fragments were obtained from Integrated DNA Technologies (IDT). Phire Hot Start II DNA Polymerase, restriction enzymes, T4 DNA ligase, and *Dpn* I were purchased from Fisher Scientific. Angiotensin I, dynorphin A, pentane, hexadecane, and Trizma base were obtained from Sigma-Aldrich. 1,2-diphytanoyl-sn-glycero-3-phosphocholine (DPhPC) was purchased from Avanti Polar Lipids. Sodium chloride and Triton X-100 was bought from Carl Roth.

### Plasmid Construction for proteins

gBlock gene fragments were ordered for synthesis by IDT, and cloned into pT7-SC1 plasmid^41^ using *Nco* I and *Hind* III restriction digestion sites. Plasmid and gene were ligated together using T4 ligase (Fermentas). 1.0 μL of the ligation mixture was incorporated into 50 µL E. cloni® 10G (Lucigen) competent cells by electroporation. Transformants were grown overnight at 37 °C on LB agar plates supplemented with ampicillin (100 μg/mL). Ampicillin-resistant colonies were picked and inoculated into 5 mL LB medium supplemented with ampicillin (100 μg/mL) for plasmid DNA preparation. The plasmid was extracted with GeneJET Extraction Kit (Fisher Scientific). The identity of the clones was confirmed by sequencing at Macrogen.

### Plasmid Construction for building a sequencing proteasome machine

gBlock gene fragments of *Thermoplasma acidophilum* α and β were ordered for synthesis by IDT. The gene encoding for the α-subunit was cloned upstream of pETDuet-1 vector (Novagen) between the *Nco* I and *Hind* III sites with the gene of Strep-tag at the C-terminus. Subsequently, the gene encoding for an untagged β-subunit was cloned downstream between the *Nde* I and *Kpn* I sites. PA-nanopore was fused to α-subunit gene through PCR splicing by overlap extension^42^, and cloned into pRSFDuet-1 vector (Novagen) using *Nco* I and *Hind* III restriction digestion sites with His tag at the N terminus.

### Construction of mutants

All mutants were constructed using the QuickChange protocol^43^ for site-directed mutagenesis on a circular plasmid template DNA with Phire Hot Start II Polymerase. Partially overlapping primers were used to avoid primer self-extension. PCR amplification was as follows: denaturation at 98 °C for 3 min, followed by 30 cycles of 98 °C for 30 s, 55°C for 30 s, and 72 °C for 2.5 min, and a final extension cycle of 72°C for 5 min. After the PCR reaction, the parental DNA template was digested with *Dpn* I enzyme for 1 h at 37°C. The PCR amplified plasmid was separated on 1% agarose gel, extracted with GeneJET Gel Extraction Kit (Fisher Scientific), and incorporated into 50 µL E. cloni® 10G (Lucigen) competent cells by electroporation. Transformants containing the plasmid were grown overnight at 37 °C on LB agar plates supplemented with ampicillin (100 μg/mL). Ampicillin-resistant colonies were picked and inoculated into 5 mL LB medium supplemented with of ampicillin (100 μg/mL) for plasmid DNA preparation. The plasmid was extracted with GeneJET Extraction Kit (Fisher Scientific), and sequenced at Macrogen for confirmation of the mutation.

### Expression and purification

The gene of the PA-nanopore was transformed into *E. coli*. BL21 (DE3) pLysS chemically competent cells. Transformants were selected after overnight growth at 37 °C on lysogeny broth (LB) agar plates supplemented with ampicillin (100 mg/L). The resulting colonies were inoculated into 200 mL LB medium containing 100 mg/L of ampicillin. The cells were grown at 37 °C (180 rpm shaking). After the optical density reached an absorbance of 0.6 at 600 nm, the expression was induced by addition of 0.5 mM isopropyl β-D-1-thiogalactopyranoside (IPTG). The temperature was lowered to 25 °C, and the cell cultures were further grown overnight. The cells were harvested by centrifugation for 20 min (4000 x g) at 4 °C and the pellets were stored at -80 °C. About 100 mL of cell culture pellet was thawed and solubilized with ∼20 mL lysis buffer (150 mM NaCl, 50 mM Tris-HCl, pH 7.5, 1 mM MgCl_2_, 0.1 units/mL DNase I, 10 µg/mL lysozyme, 1% v/v Triton X-100) and stirred with a vortex shaker for 1 hour at 22 °C. The bacteria were then lysed by sonication (duty cycle 10%, output control 3, Branson Sonifier 450). The lysate was subsequently centrifuged at 6000 x g at 4 °C for 20 min and the cellular debris discarded. The supernatant was mixed with 100 μL of Strep-Tactin resin (IBA) to a 50 mL falcon tube, which was pre-equilibrated with wash buffer (1% v/v Triton X-100, 150 mM NaCl, 15 mM Tris-HCl, pH 7.5). After 1 hour, the resin was loaded into a column (Micro Bio Spin, Bio-Rad), which was pre-washed with 5 mL wash buffer (150 mM NaCl, 50 mM Tris-HCl, pH 7.5, 1% v/v Triton X-100). In total, 10 mL of wash buffer (0.5% v/v Triton X-100, 150 mM NaCl, 50 mM Tris, pH 7.5, 20 mM imidazole) was used to wash the beads. The protein was eluted with approximately 100 μL elution buffer (2.5 mM desthiobiotin, 150 mM NaCl, 50 mM Tris-HCl, pH 7.5, 0.5% v/v Triton X-100).

The gene encoding for S1, VATΔN, and GFP were separately transformed into *E. coli*. BL21 (DE3) electrocompetent cells. Transformants were selected after overnight growth at 37 °C on lysogeny broth (LB) agar plates supplemented with ampicillin (100 mg/L). The resulting colonies were inoculated into 200 mL LB medium containing 100 mg/L of ampicillin. The cells were grown at 37°C (180 rpm shaking). After the optical density reached an absorbance of 0.6 at 600 nm, the expression was induced by addition of 0.5 mM isopropyl β-D-1-thiogalactopyranoside (IPTG) at 25 °C. And the cell cultures were further grown overnight. The cells were harvested by centrifugation for 20 min (4000 x g) at 4 °C and the pellets were stored at -80 °C. About 100 mL of cell culture pellet was thawed and solubilized with ∼20 mL lysis buffer (150 mM NaCl, 50 mM Tris-HCl, pH 7.5, 1 mM MgCl_2_, 0.1 units/mL DNase I, 10 µg/mL lysozyme) and stirred with a vortex shaker for 1 hour at 4°C. The bacteria were then lysed by sonication (duty cycle 10%, output control 3, Branson Sonifier 450). The lysate was subsequently centrifuged at 6000 x g at 4 °C for 20 min and the cellular debris discarded. The supernatant was mixed with 100 μL of Ni-NTA resin (Qiagen) to a 50 mL falcon tube, which was pre-equilibrated with wash buffer (150 mM NaCl, 50 mM Tris-HCl, pH 7.5). After 1 hour at 4 °C, the resin was loaded into a column (Micro Bio Spin, Bio-Rad), which was pre-washed with 5 mL wash buffer (150 mM NaCl, 50 mM Tris-HCl, pH 7.5). In total, 10 mL of wash buffer (150 mM NaCl, 50 mM Tris, pH 7.5, 20 mM imidazole) was used to wash the beads. The protein was eluted with approximately 200 μL elution buffer (500 mM imidazole, 150 mM NaCl, 50 mM Tris-HCl, pH 7.5).

### Proteasome co-expression and purification

For the assembly of the proteasome-nanopore, the pETDuet-1 containing the gene encoding for the α and β-subunits of the proteasome and pRSFDuet-1 containing the gene encoding for the PA28-αΔ20 nanopore plasmids were co-transformed into *E. coli* BL21 (DE3) electrocompetent cells. Transformants were selected after overnight growth at 37 °C on LB agar plates supplemented with ampicillin (100 mg/L) and kanamycin (100 mg/L). The resulting colonies were inoculated into 200 mL LB medium containing 100 mg/L of ampicillin and kanamycin. Protein expression was induced by 0.5 mM β-d-thiogalactopyranoside (IPTG) when the A600 reached about 0.6. The temperature was lowered to 25°C. After 12 h induction, the cells were collected, and the pellets were stored at -80 °C.

About 100 mL of cell culture pellet was thawed and solubilized with ∼20 mL lysis buffer (150-1000 mM NaCl, 50 mM Tris-HCl, pH 7.5, 1 mM MgCl2, 20 mM imidazole, 0.1 units/mL DNase I, 10 µg/mL lysozyme, 1% v/v Triton X-100) and stirred with a vortex shaker for 1 hour at 22 °C. The bacteria were then lysed by sonication (duty cycle 10%, output control 3, Branson Sonifier 450). The lysate was subsequently centrifuged at 6000 x g at 4 °C for 20 min and the cellular debris discarded. The supernatant was mixed with 100 μL of Ni-NTA resin (Qiagen) to a 50 mL falcon tube, which was pre-equilibrated with wash buffer (1% v/v Triton X-100, 150 mM NaCl, 50 mM Tris-HCl, pH 7.5). After 1 hour, the resin was loaded into a column (Micro Bio Spin, Bio-Rad), which was pre-washed with 5 mL wash buffer (150 mM NaCl, 15 mM Tris-HCl, pH 7.5, 1% v/v Triton X-100). The protein was eluted with approximately 200 μL elution buffer (500 mM imidazole, 150-1000 mM NaCl, 15 mM Tris-HCl, pH 7.5, 1% v/v Triton X-100). Subsequently, the eluted protein was mixed with 50 μL of Strep-Tactin resin (IBA) to a 2 mL tube, which was pre-equilibrated with wash buffer (1% v/v Triton X-100, 150 mM NaCl, 15 mM Tris-HCl, pH 7.5). After 30 minutes, the resin was loaded into a column (Micro Bio Spin, Bio-Rad), which was pre-washed with 5 mL wash buffer (150 mM NaCl, 50 mM Tris-HCl, pH 7.5, 1% v/v Triton X-100). In total, 10 mL of wash buffer (150-1000 mM NaCl, 50 mM Tris, pH 7.5, 20 mM imidazole, 0.5% v/v Triton X-100) was used to wash the beads. The protein was eluted with approximately 100 μL elution buffer (2.5 mM desthiobiotin, 150-1000 mM NaCl, 50 mM Tris-HCl, pH 7.5, 0.5% v/v Triton X-100).

### Proteolytic activity of artificial proteasome-nanopore

To determine the proteolytic activity of artificial proteasome-nanopore, β-casein was incubated with purified proteasome-nanopore (complex 3, Fig. 3) under a variety of incubating time, temperature, and salt concentration. Firstly, an aliquot of 0.1 mL β-casein (1 mg/mL) was incubated with complex 3 at 53 °C in buffer A (50 mM Tris, pH 7.5, 150 mM NaCl). The final β-casein/ proteasome-nanopore concentration ratio was 42. In the absence of the protease, no degradation of β-casein was observed. After 15 min of incubation at 53 °C with the proteasome-nanopore, almost all β-casein was digested, with about three quarters of the initially observed proteins no longer detectable on SDS-PAGE. After 30 minutes’ incubation, all β-casein was digested. Then, a variety of temperature and salt concentration for degradation of β-casein were tested. The proteolytic activity increased with the temperature and decreased with increasing the salt concentration.

### Electrical recordings in planar lipid bilayers

The setup consisted of two chambers separated by a 25 μm thick polytetrafluoroethylene film (Goodfellow Cambridge Limited), which contain an aperture of approximately 100 μm in diameter, which was formed by applying a high voltage spark. To form a lipid bilayer, the aperture was pre-treated with a drop of 5% hexadecane/pentane solution. After waiting about 1-5 minutes in order to allow pentane to evaporate, 500 μL of a buffered solution (150 mM NaCl, 15 mM Tris-HCl, pH 7.5) was added to each compartment. Then a drop of DPhPC in pentane (∼6 mg/mL) was added to each compartment. After evaporation of the pentane, a lipid monolayer formed spontaneously by pipetting the solution up and down over the aperture. Silver/silver-chloride electrodes were submerged into the solution of each compartment. Nanopores were added to *trans* side. All experiments were performed at ∼23 °C^44^.

### Data recordings and analysis

Electronic signals were recorded by using an Axopatch 200B (Axon Instruments) with digitization performed with a Digidata 1440 (Axon Instruments). Clampex 10.7 software and Clampfit 10.7 software (Molecular Devices) were used for electronic signal recording and subsequent data analysis, respectively. Events were collected using the single-channel search feature in Clampfit, combined together and fitted with a single exponential regression.

### Ion selectivity

The current–voltage (*I-V*) current traces were recorded with an automated voltage protocol that applied each potential for 0.22 s from -30 to +30 mV with 1 mV steps. Ag/AgCl electrodes were surrounded with 2.5% agarose bridges containing 2.5 M NaCl. Reversal potential was measured from extrapolation from *I-V* curves collected under asymmetric salt concentration condition. The experiment proceeded as follow: First an individual nanopore was reconstituted using the same buffer in both chambers (1 M NaCl, 15 mM Tris, pH 7.5, 500 µL). This allowed assessing the orientation of the nanopore and allowed balancing the electrodes. Then 500 µL solution containing 4 M NaCl, 15 mM Tris, pH 7.5 was slowly added to *trans* side and 500 µL of a buffered solution containing no NaCl (15 mM Tris, pH 7.5) was added to *cis* side (*trans*:*cis*, 2.0 M NaCl: 0.5 M NaCl).

### Transmission Electron Microscopy (TEM)

TEM experiments were carried out as following: Liquid samples were placed on carbon-coated copper grids and excess solution removed with filter paper, and then, proteins were stained using 2% uranyl acetate for 1 min. Transmission electron micrographs were imaged at 120 kV through a FEI Tecnai T20 electron microscope.

### Modelling of hybrid-nanopore via Multiscale Molecular Dynamics simulations and Protein-Protein Docking

Initially, the structures of mouse PA28α (PDB ID: 5MSJ), serine-serine-glycine linker (modelled with Pymol in straight coil configuration) and the transmembrane region of anthrax protective antigen (PDB ID: 3J9C) were aligned according to the position of the expected peptidic bonds. A Martini coarse-grain (CG) model was generated based on this initial step using Martinize^45^ and the hybrid protein was placed in a 1,2-dioleoyl-sn-glycero-3-phosphocholine (DOPC) membrane (680 lipids) using Insane. The structure was relaxed for 10 µs MD simulations. DOPC should mimic the DPhPC quite well with respect to membrane thickness (38.5/36.0 nm respectively)^46^, as well as its phase (liquid-crystalline). The last configuration of the CG MD simulation was further refined with 300 ns of all-atom (AA) simulations using the CHARMM36m force-field^47^. All CG and AA simulations were performed with the program package Gromacs (version 2018.1)^47^. All simulations were performed at 310 K, 150 mM NaCl and neutral net charge. The final configuration of the AA simulation containing the PA-nanopore and the membrane was aligned with the proteasome (PDB: 1YA7) using their PA subunits and the multiseq plug of VMD^48,49^. The nanopore was connected to the α-proteasome subunit with the linker (KGMIY) using the modeller automodel class. To finalize the model, VAT was docked on top of the nanopore complex using HADDOCK.

#### CG simulation protocols and parameters

The system was energy minimized using the steepest descent minimizer for 1000 steps). The minimized structured was equilibrated with a constrained protein backbone for 250 ps, up to the point that the β-barrel was fully hydrated (dt = 1 fs). To relax the protein in the membrane one more unconstrained simulation at equal time step was performed for another 250 ps. CG production MD proceeded 10 µs (dt = 10 fs). The CG simulations were coupled to a v-rescale thermostat^50^ set at 310 K at 1 ps interval, water/ions; protein; and lipids were coupled independently. Pressure was maintained at 1 bar using a semi-isotropic Berendsen^50^ (3 ps interval) and Parrinnello-Rahman (12 ps interval) barostat^51^ for equilibration and production, respectively. Electrostatics and Lennard-Jones were handled using the default Martini^52^ settings (0 nm potential-shit-verlet, 1.1 nm reaction field; and 0 nm potential-shit-verlet, 1.1 nm cutoff respectively).

#### Backmapping and AA simulation protocols and parameters

The whole system from the final frame in the CG simulation (10 µs) was backmapped to the CHARMM36m force field using the default minimization and equilibration scheme described by Wassenaar *et al*.^53^ (DOPC → DPhPC). After equilibration, a production run was performed spanning 300 ns. The default CHARMM36m settings were used for the time step (2 fs), Lennard-Jones (1.0 nm force-switch; 1.2 nm cut-off) and coulombic interactions (pme; 1.2 nm potential-shift-verlet). Semi-isotropic pressure coupling was performed using the Parrinello-Rahman barostat^51^ (1 Bar at 5 ps intervals) and the Nose-Hoover^54^ thermostat (1 ps intervals) was used to maintain a constant temperature of 310 K.

#### HADDOCK protocol and parameters

Docking was performed using HADDOCK2.4^55^. The VAT structure (PDB: 1YA7) and the α-subunits of 20S proteasome (PDB: 5G4G) were used for docking. We defined E714 and K718 of VAT and K20 and E29 of the α-subunits as active residues, with the passive residues defined with a cutoff of 4.5 Å. These residues were considered important for the alignment of the pore, as they should be involved in complementary salt-bridges between the complexes in the protein-protein interface. The default settings suggested by the web-server interface^55^ were used for the docking. The two best clusters found show a small misalignment of the complexes, which is typical for VAT-proteasome interactions, as there is a mismatch in the number of subunits (6 monomers in VAT and 7 α-subunits of 20S proteasome)^56^. Cluster one was selected, as showed the most favorable VdW and electrostatic interactions. The pores of the VAT and the α-subunits were aligned as expected for such a complex, showing a straight uninterrupted path through the whole complex.

**Supplementary Fig. 1.**
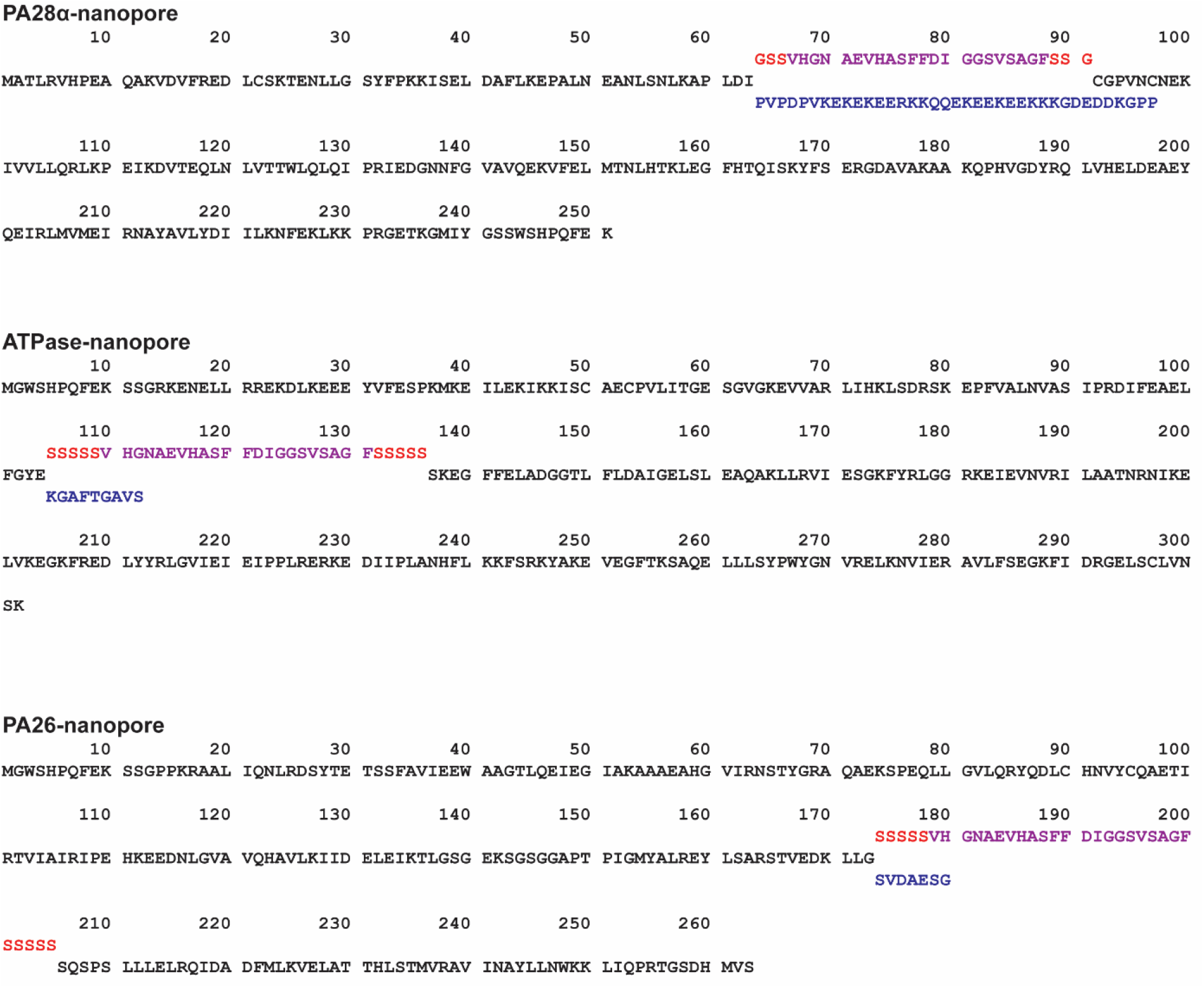
Complete sequences of artificial-nanopore. In black and blue is the sequence of soluble proteins, in red (linker) and purple (transmembrane region of the protective antigen) the inserted polypeptide.

**Supplementary Fig. 2.**
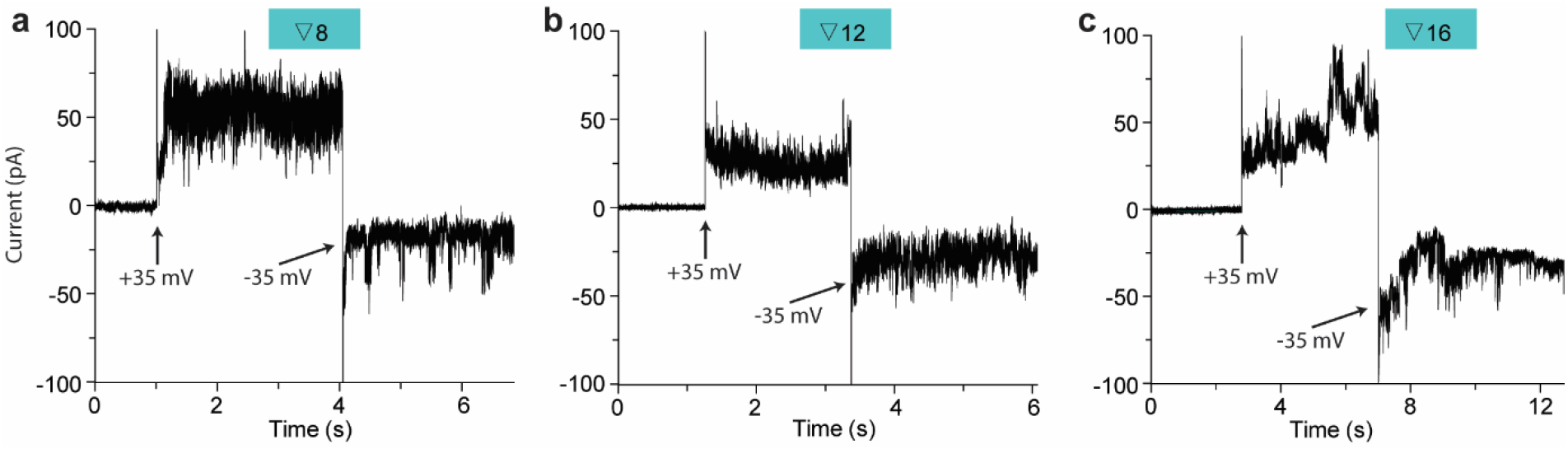
Electrical properties of insertion mutants at ±35 mV. Electrical recordings of a single nanopore at ±35 mV. Data were collected at ±35 mV in 1 M NaCl, 15 mM Tris, pH 7.5, using 10 kHz sampling rate and a 2 kHz low-pass Bessel filter.

**Supplementary Fig. 3.**
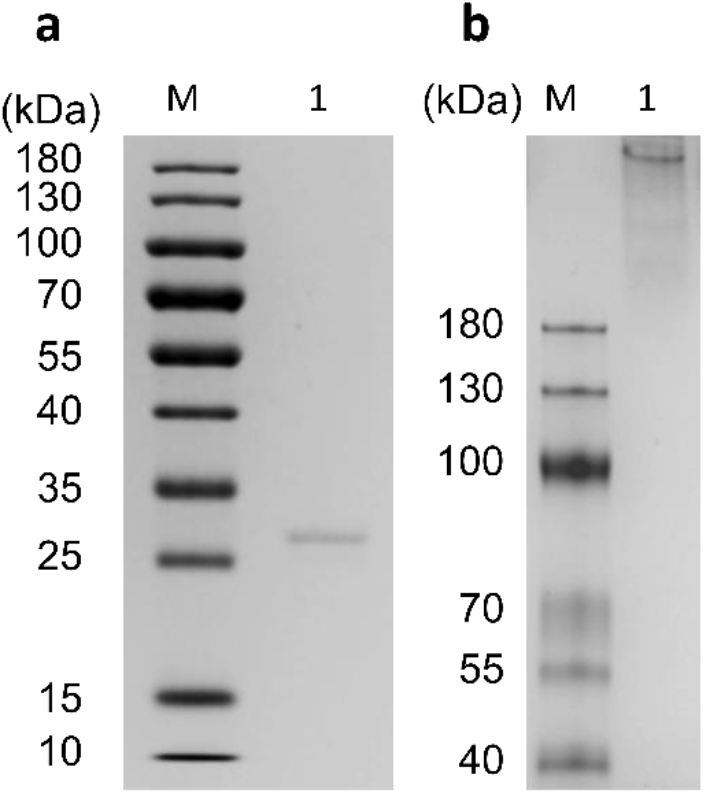
SDS-PAGE (a) and Native PAGE (b) analyses of artificial nanopore. Lane 1, ∇2 mutant (∼1µg); Lane M, protein markers and their corresponding molecular masses.

**Supplementary Fig. 4.**
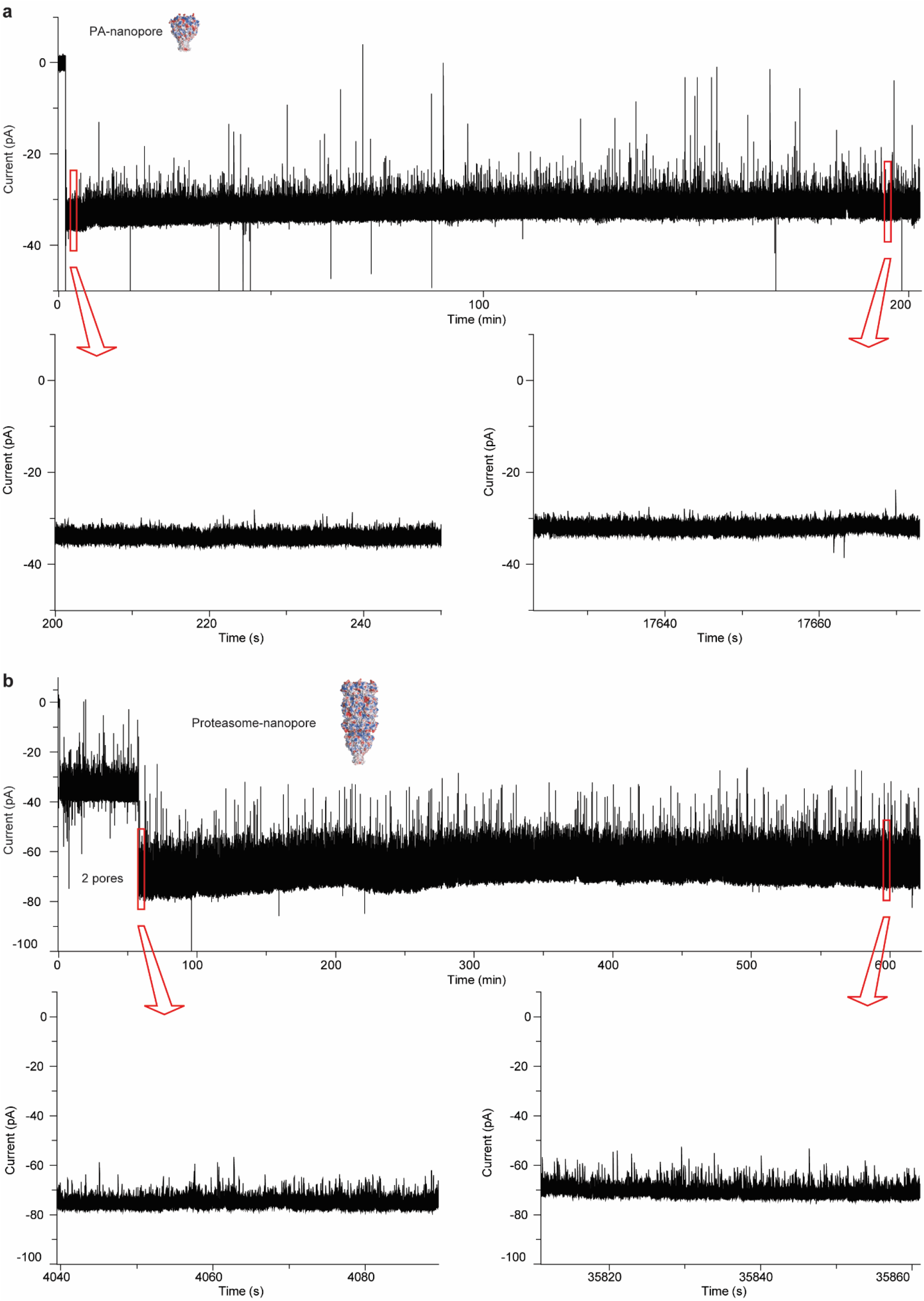
Continuous current trace recording of the PA-nanopore and proteasome-nanopore. Data were collected at 22 °C and -30 mV in 1 M NaCl, 15 mM Tris, pH 7.5, using a 2 kHz low-pass Bessel filter with a 10 kHz sampling rate.

**Supplementary Fig. 5.**
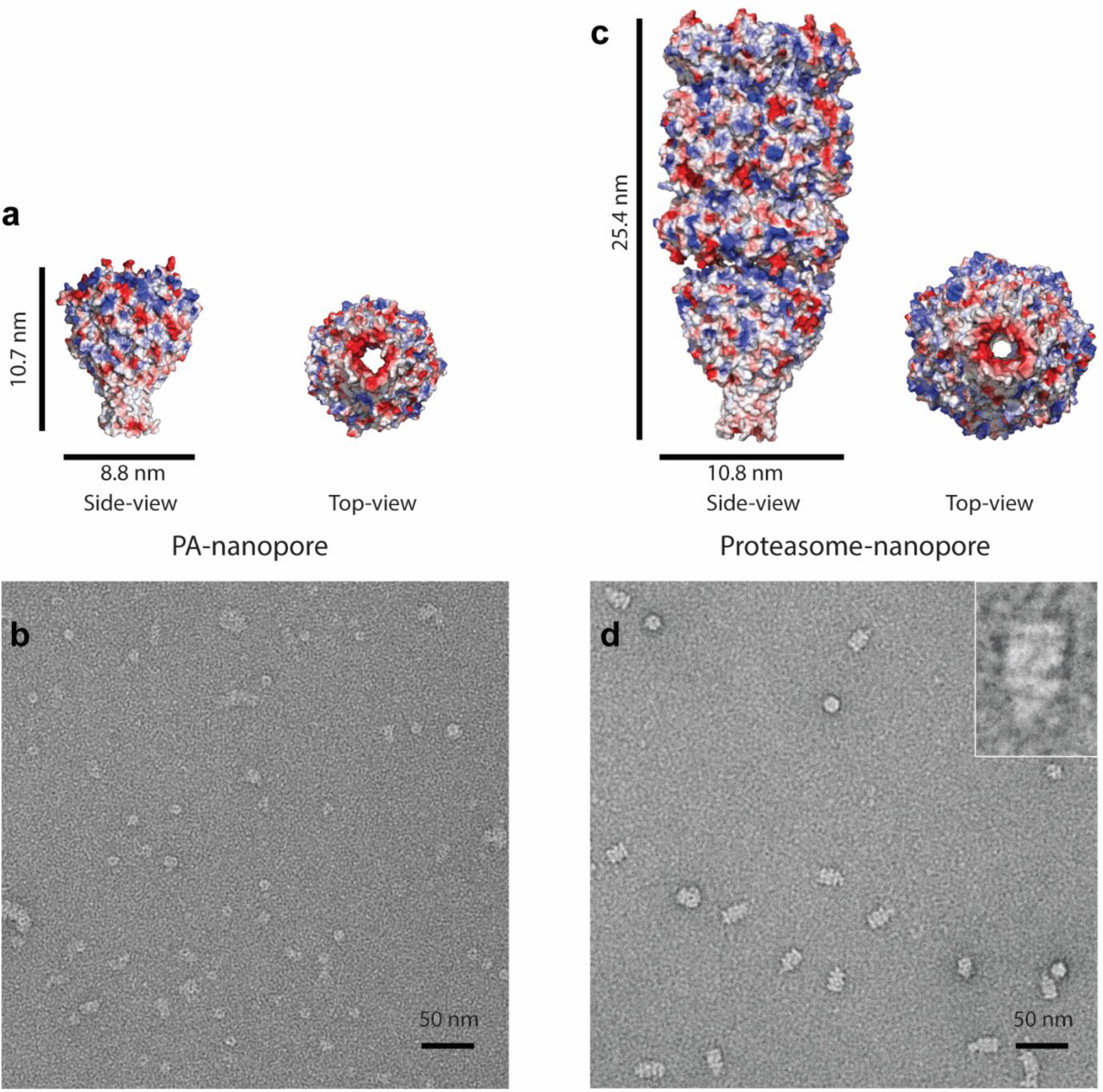
Surface representation and negatively stained TEM images. **a**, and **c**, surface representations of the PA-nanopore and the proteasome-nanopore coloured according to the vacuum electrostatic potential as calculated by PyMOL on the final snapshot of the multiscale MD model. **b** and **d**, The PA-nanopore and proteasome-nanopore were negatively stained with 2% uranyl acetate.

**Supplementary Fig. 6.**
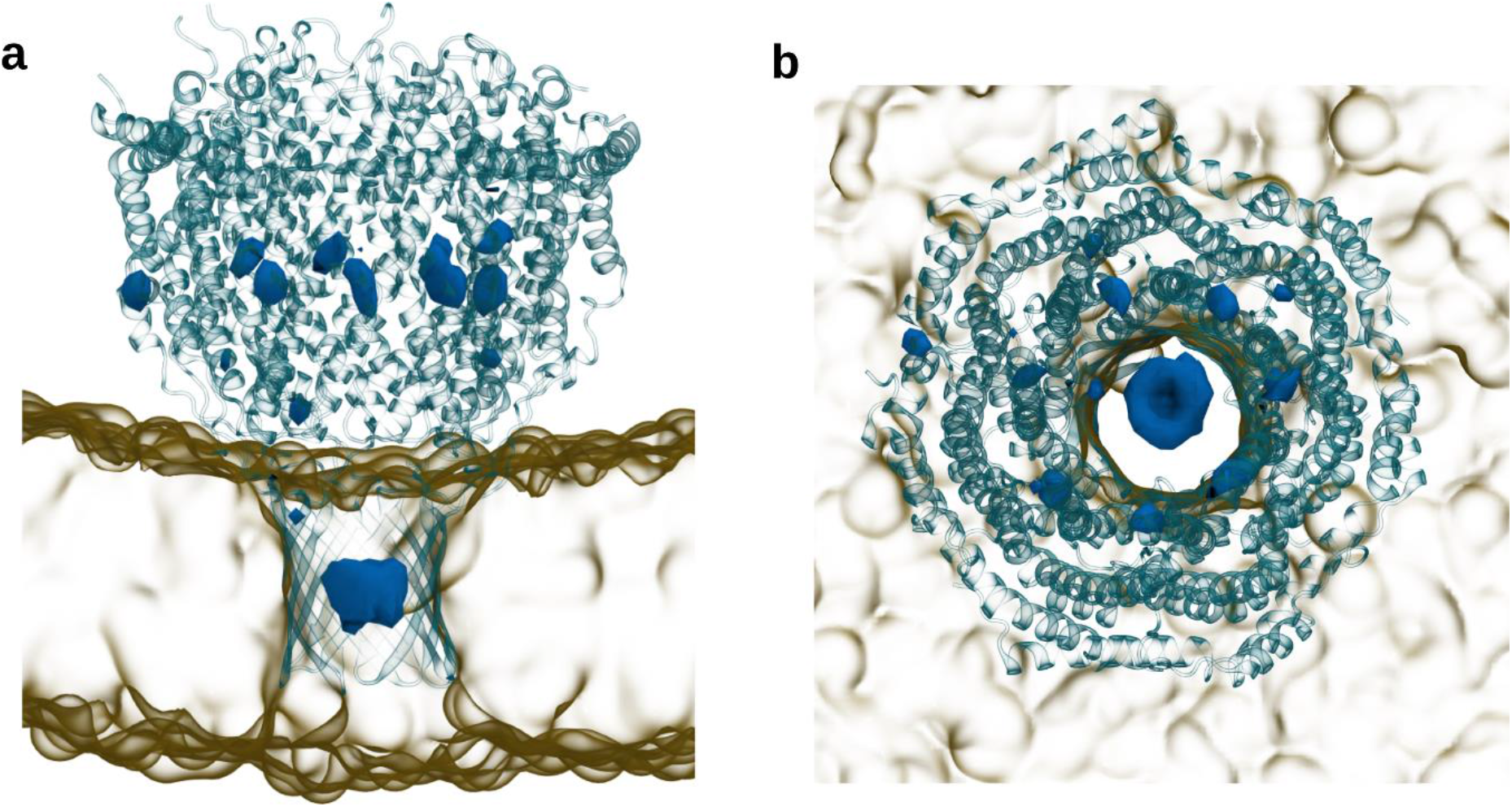
Sodium ion density within the hybrid artificial nanopore obtained from averaging ∼300 ns atomistic simulations. The densities shown in blue volume maps correspond to an isovalue of at least 10 times the bulk density, computed with VMD Volume map plugin. There are two main regions which we observe higher occupancy of sodium ions: (i) between residues GLU 302, GLU308 and ASP315 of each monomer of the transmembrane beta-barrel of the protective antigen; (ii) near to GLU168 and GLU211 of PA28α unit of the proteosome. There is no meaningful density of chloride ions inside the channel.

**Supplementary Fig. 7.**
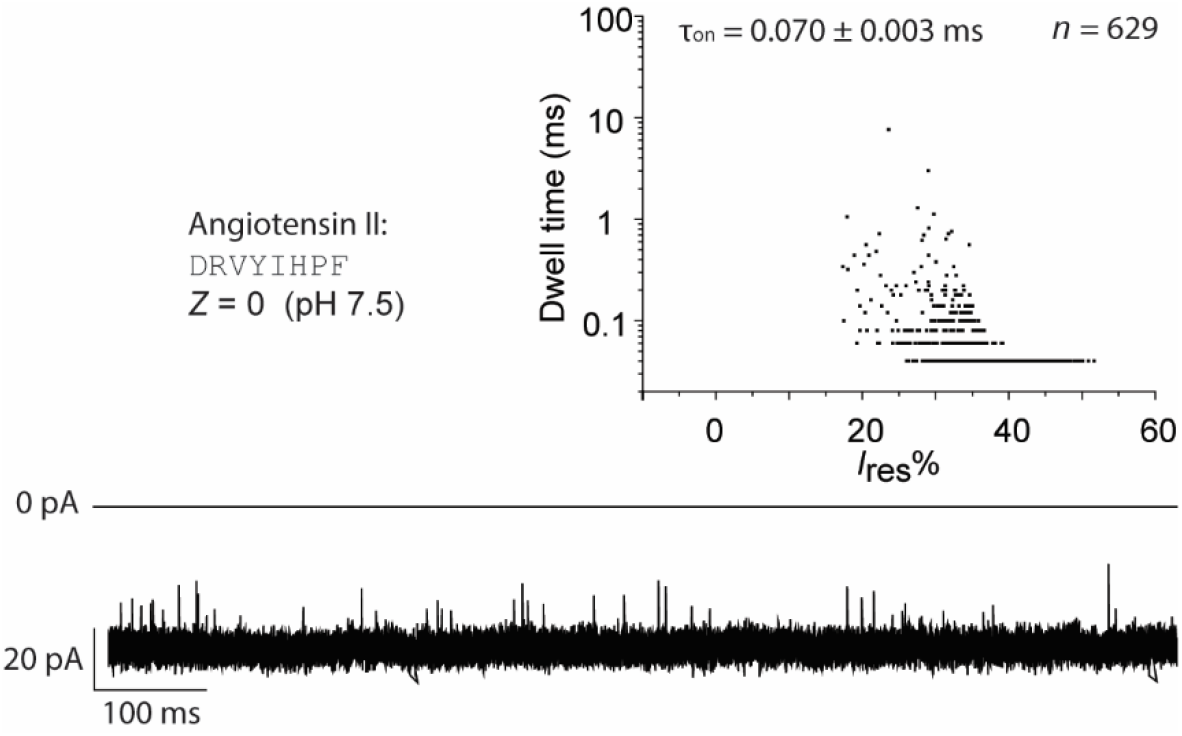
The capture of angiotensin II by PA-nanopre. Peptide sequences and net charge of angiotensin II (left), scatter plots of *I*_res_% versus dwell time (right), and representative trace (below). The current signals were filtered at 10 kHz and sampled at 50 kHz.

**Supplementary Fig. 8.**
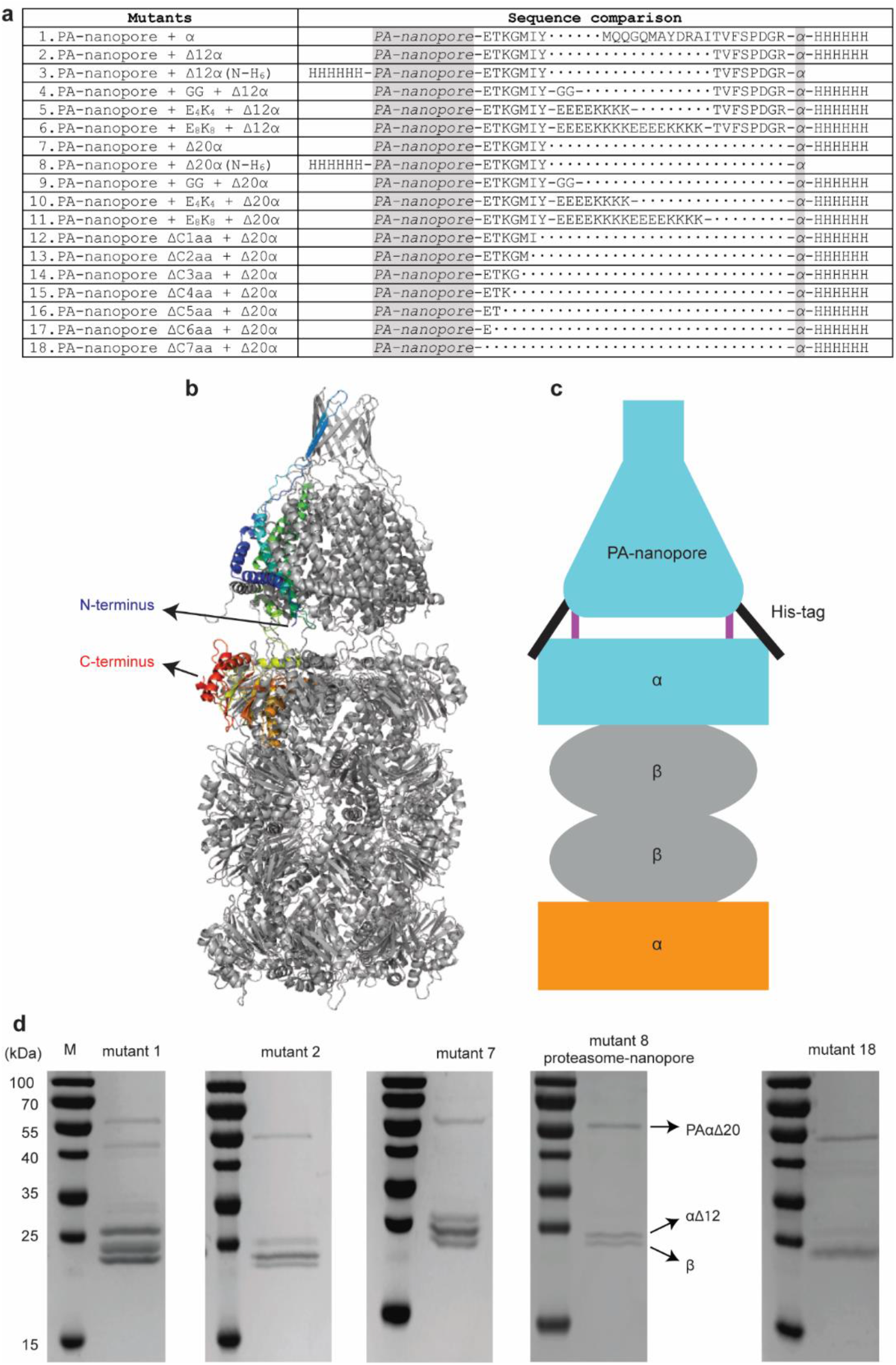
Optimizing proteasome-nanopore. **a**, Mutants list tested in this work. **b**, Cartoon (left) representation of the proteasome-nanopore. One subunit of fusion protein (PA-nanopore + α) was coloured progressively from blue (N-terminus) to red (C-terminus). **c**, Schematic representation of the proteasome-nanopore showing a speculative mechanism leading to the protection of the linker from proteolytic cleavage, in which the N-terminal His-tags shield the groove between PA-nanopore and α. **d**, SDS-PAGE analyses of the purified proteasome-nanopore mutants. After co-expression with β-subunit and a second α-subunit, only mutant 8 (proteasome-nanopore) presents three unique bands.

**Supplementary Fig. 9.**
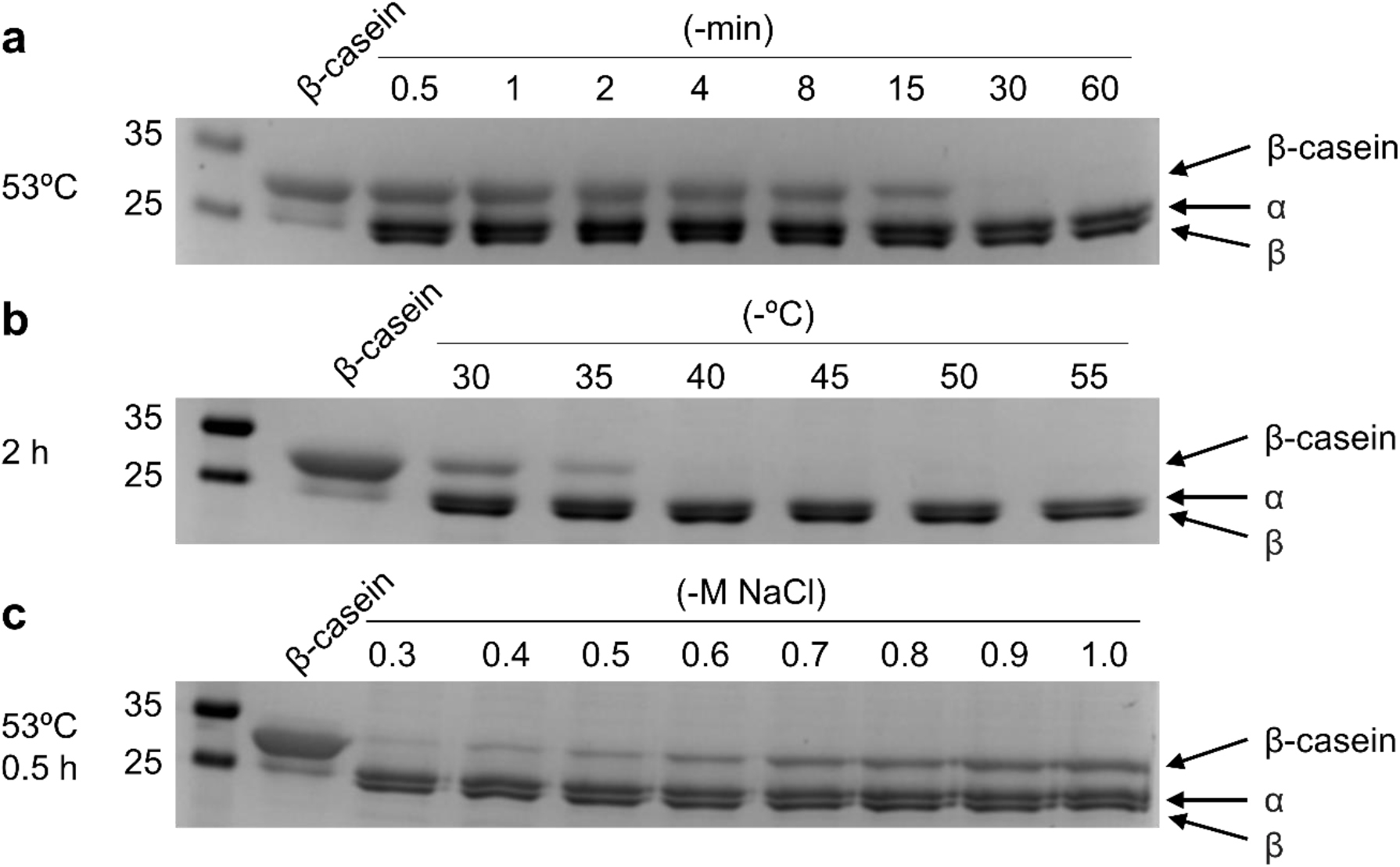
SDS-PAGE analysis the hydrolyzing activity of proteasome-nanopore. **a**, β-casein (1 mg/mL) was incubated with subcomplex 3 at 53 °C in buffer A (50 mM Tris, pH 7.5, 150 mM NaCl). **b**, β-casein (1 mg/mL) was incubated with proteasome-nanopore for 2 hours in buffer A. **c**, β-casein (1 mg/mL) was incubated with proteasome-nanopore at 53 °C for 0.5 hour in buffer B (50 mM Tris, pH 7.5, 0.3-1.0 M NaCl). The β-casein/proteasome-nanopore concentration ratio was 42.

**Supplementary Fig. 10.**
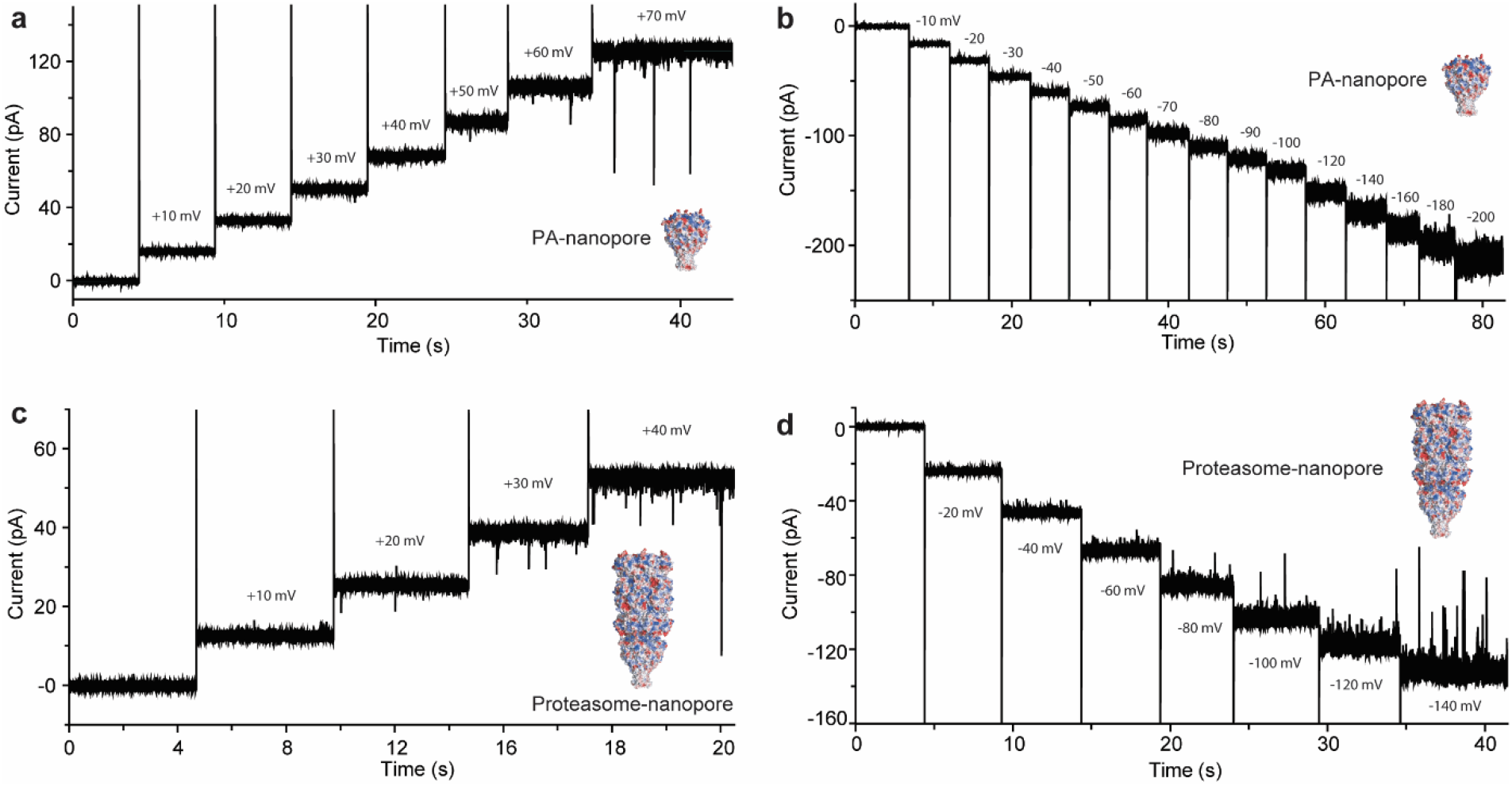
The stability of the PA-nanopore and proteasome-nanopore. **a** and **b**. PA-nanopore shows gating above approximately +50 mV, but is stable at potentials as high as -200 mV. **c** and **d**. Proteasome-nanopore displays gating above potentials of approximately +20 mV and -80 mV. Data were collected at 22 °C in 1 M NaCl, 15 mM Tris, pH 7.5, using a 2 kHz low-pass Bessel filter with a 10 kHz sampling rate.

**Supplementary Fig. 11.**
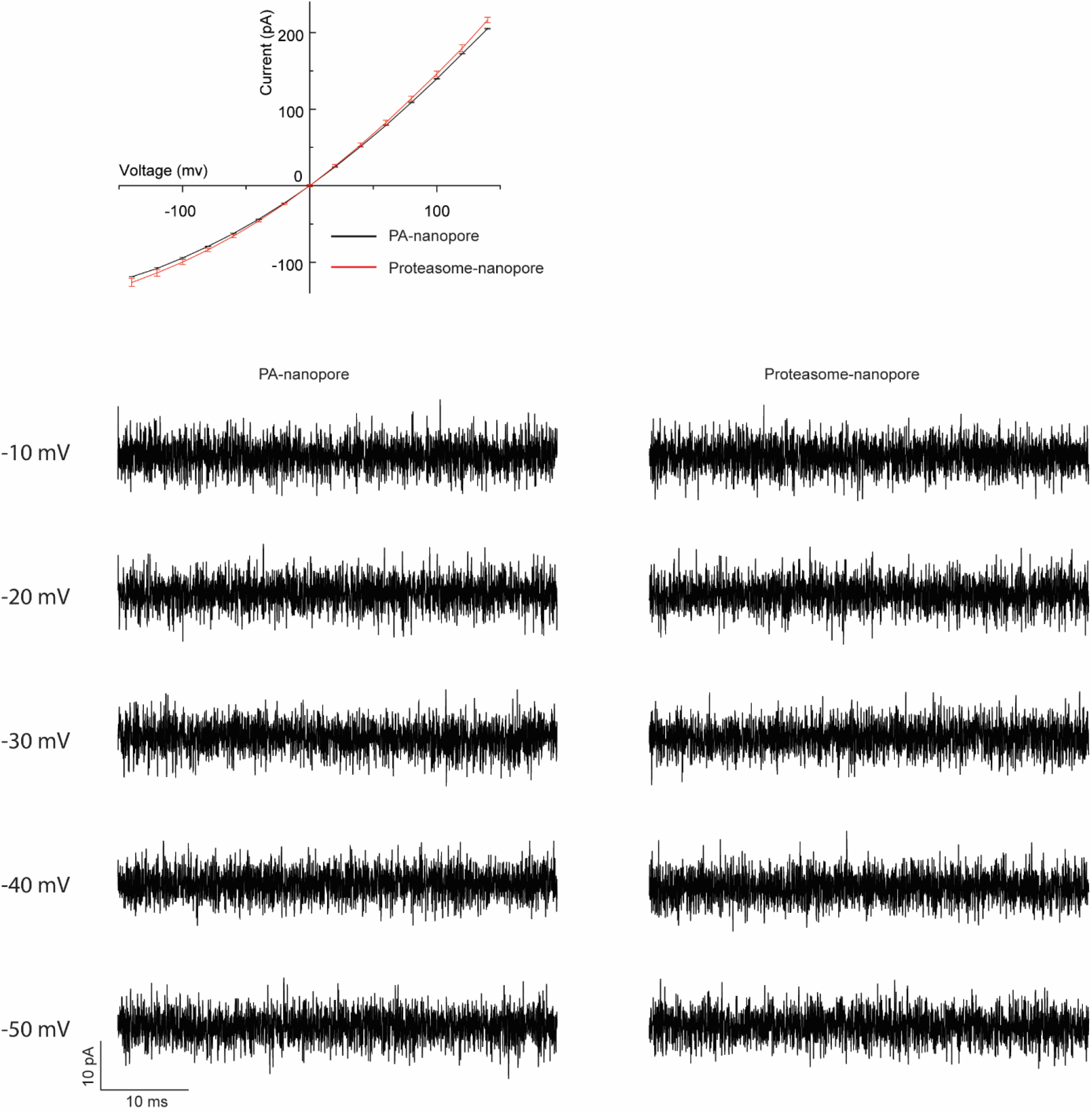
Averaged current–voltage (*I*–*V*) characteristics of three different nanopores (top) and noise comparison (bottom). The error bars represent one standard deviation. The artificial transmembrane proteasome did not alter the conductance of the nanopore. The current signals were recorded at -10, -20, -30, -40 and -50 mV in 1 M NaCl, 15 mM Tris, pH 7.5, filtered at 10 kHz, and sampled at 50 kHz.

**Supplementary Fig. 12.**
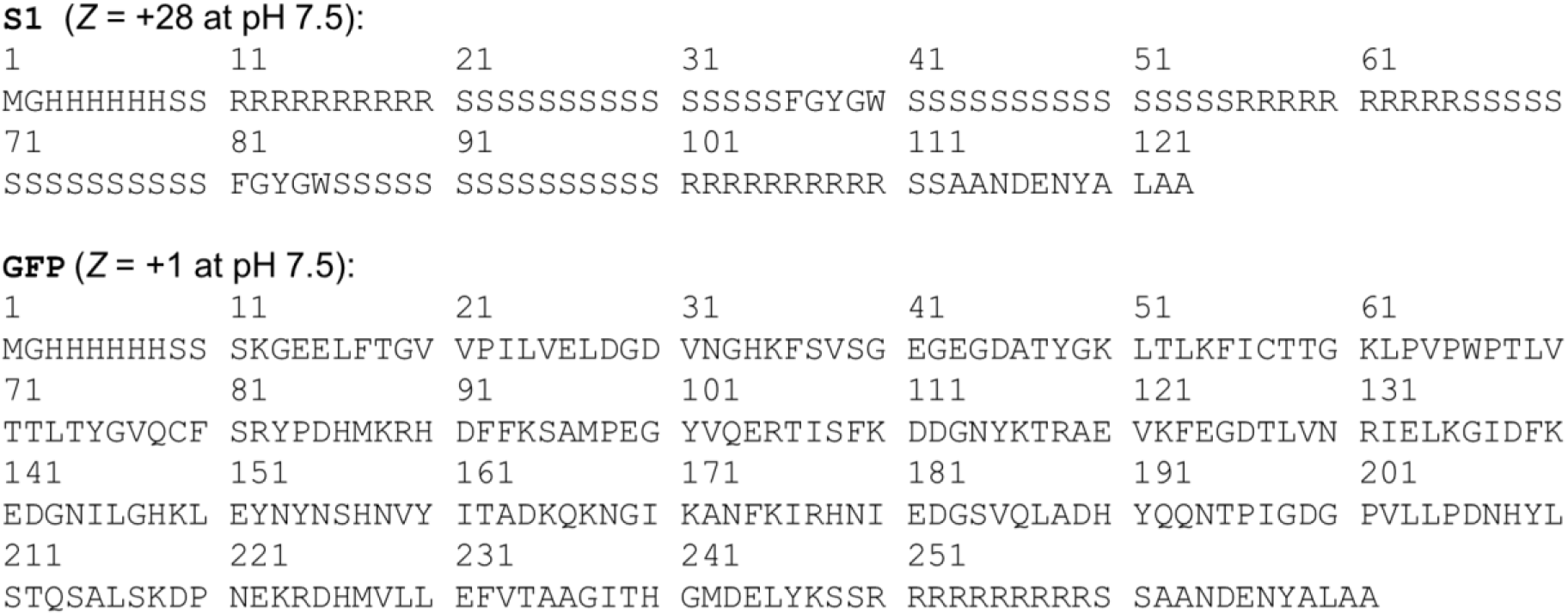
Primary sequences of S1, ssrA-tagged GFP.

**Supplementary Fig. 13.**
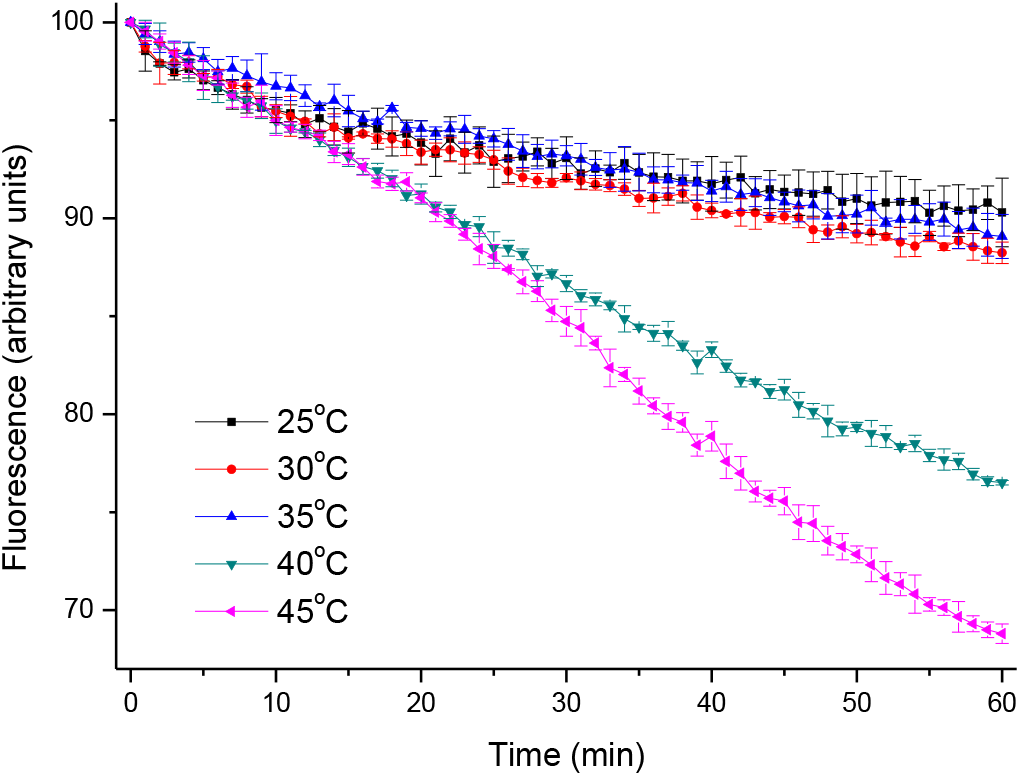
Bulk phase assays of degradation of superfolded GFP bearing an ssrA tag at various temperatures. The time course of fluorescence changes of 2 µM GFP in the presence of 50 nM VATΔN was followed at various temperatures in 0.1 M PBS buffer, pH 7.5 (including 20 mM MgCl_2_, 50 nM proteasome, and 5 mM ATP).

**Supplementary Fig. 14.**
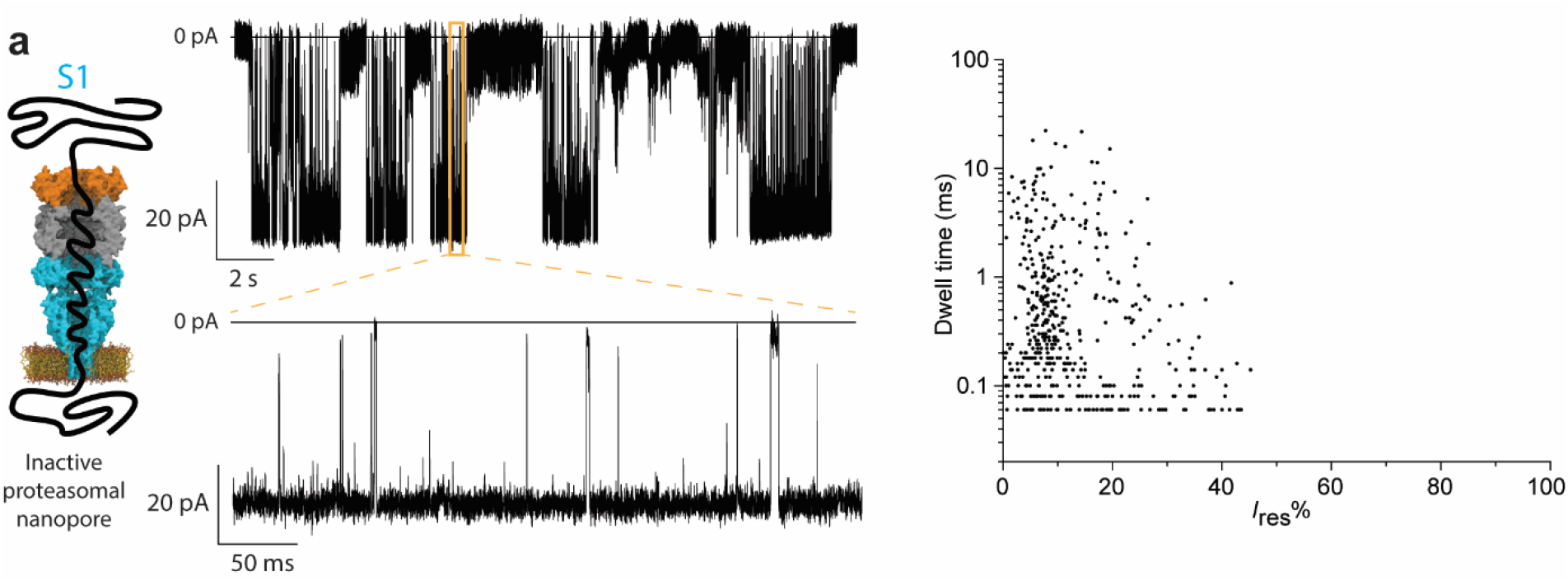
Typical current trace provoked by substrate 1 (S1) using an inactive proteasome-nanopore.

**Supplementary Fig. 15.**
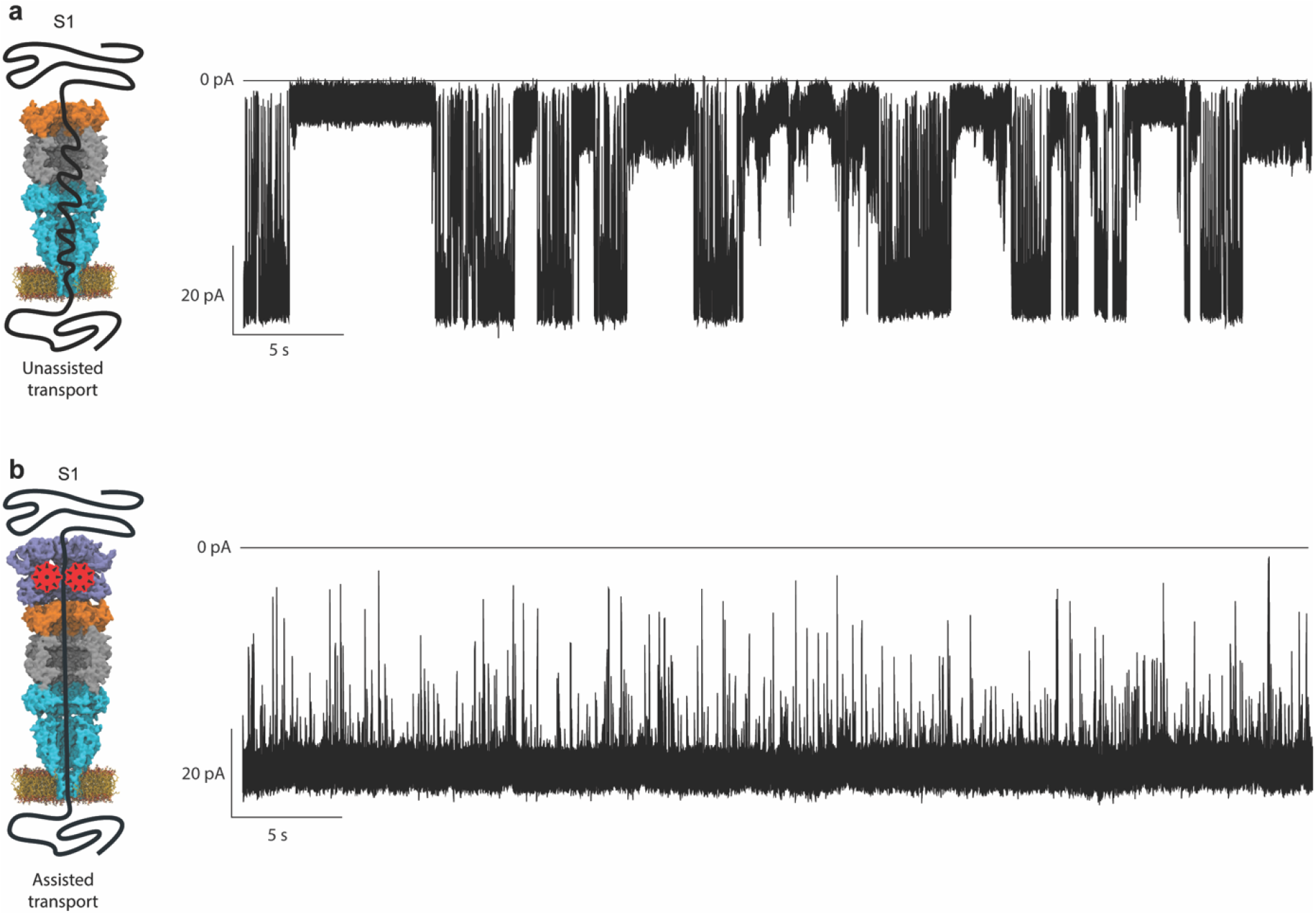
Assisted S1 translocation. **a**, translocation of S1 through n inactivated proteasome-nanopore. **b**, translocation of S1 mediated by VATΔN. The proteasome-nanopore and substrates were added to the *cis* side. Data were collected at 40 °C and -30 mV in 1 M NaCl, 15 mM Tris, pH 7.5, using a 10 kHz low-pass Bessel filter with a 50 kHz sampling rate. The traces were then filtered digitally with a Gaussian low-pass filter with a 5 kHz cut-off.

**Supplementary Fig. 16.**
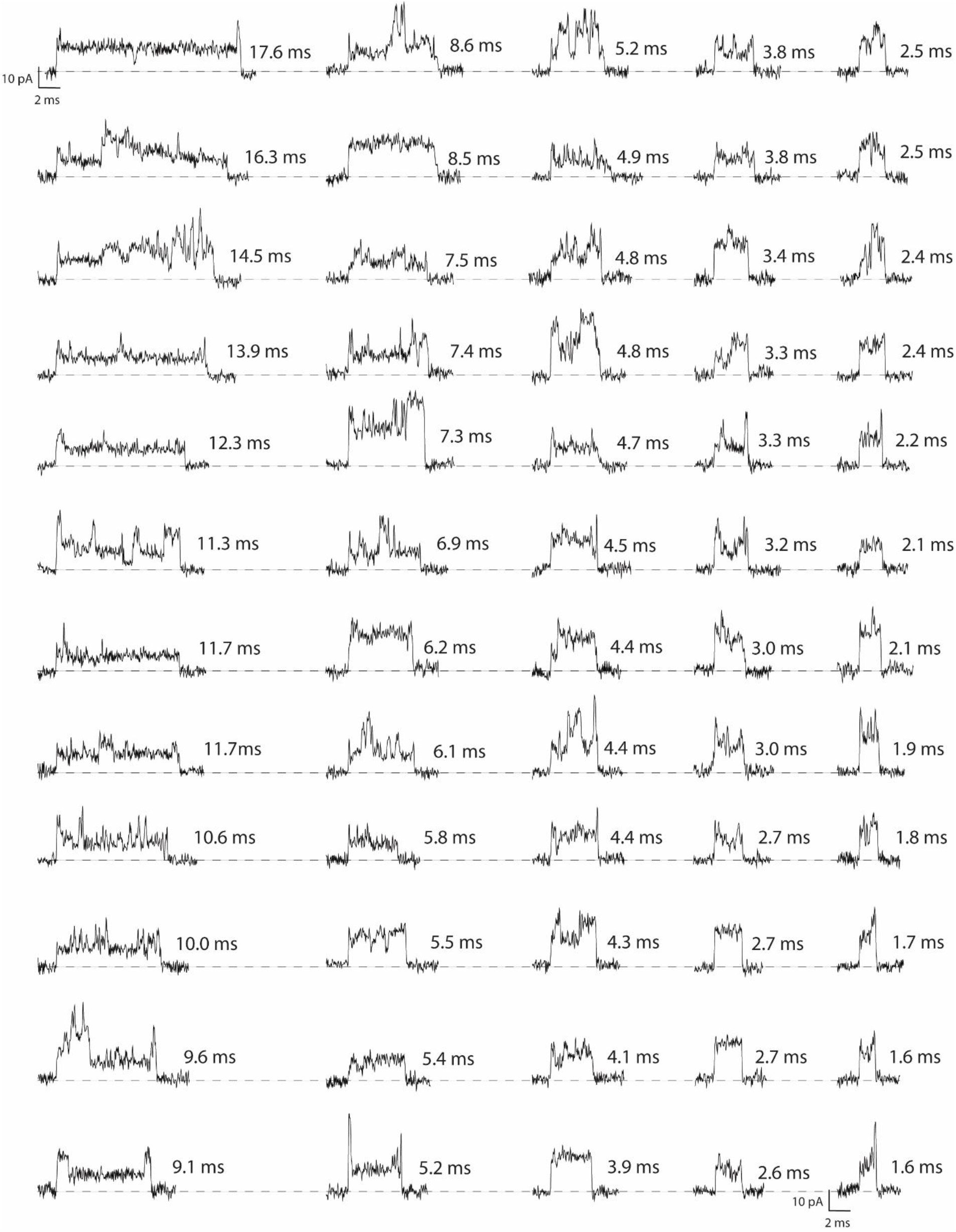
Representative blockades of the S1 translocation though inactivated proteasome-nanopore in the presence of 5 µM VATΔN at 40 °C in 1 M NaCl, 15 mM Tris, pH 7.5. The current signals were recorded at -30 mV, filtered at 10 kHz, and sampled at 50 kHz. The traces were then filtered digitally with a Gaussian low-pass filter with a 5 kHz cut-off.

**Supplementary Fig. 17.**
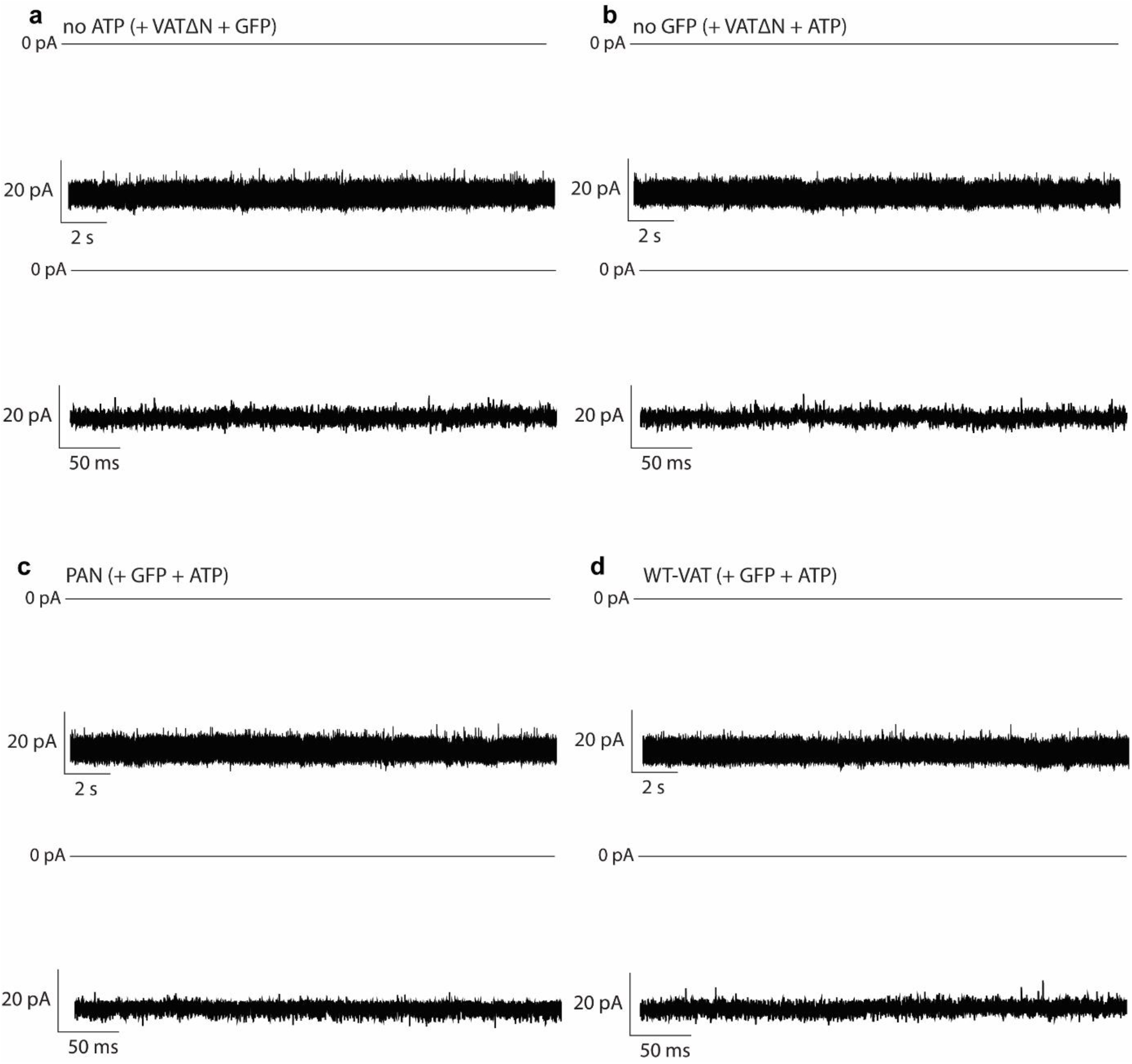
Representative traces of control experiments. **a**, Representative traces in the absence of ATP but in the presence of 5 µM VATΔN and 5µM GFP. **b**, Representative traces in which no GFP was used but in the presence of 5 µM VATΔN and 2 µM ATP. **c** and **d**, Representative traces in which PAN or WT-VAT were used instead of VATΔN in the presence of 2 µM ATP and 5 µM GFP. Data were collected at -30 mV and 40 °C, in 1 M NaCl, 15 mM Tris, pH 7.5. The current signals were filtered at 10 kHz, and sampled at 50 kHz. The traces were then filtered digitally with a Gaussian low-pass filter with a 5 kHz cut-off.

**Supplementary Fig. 18.**
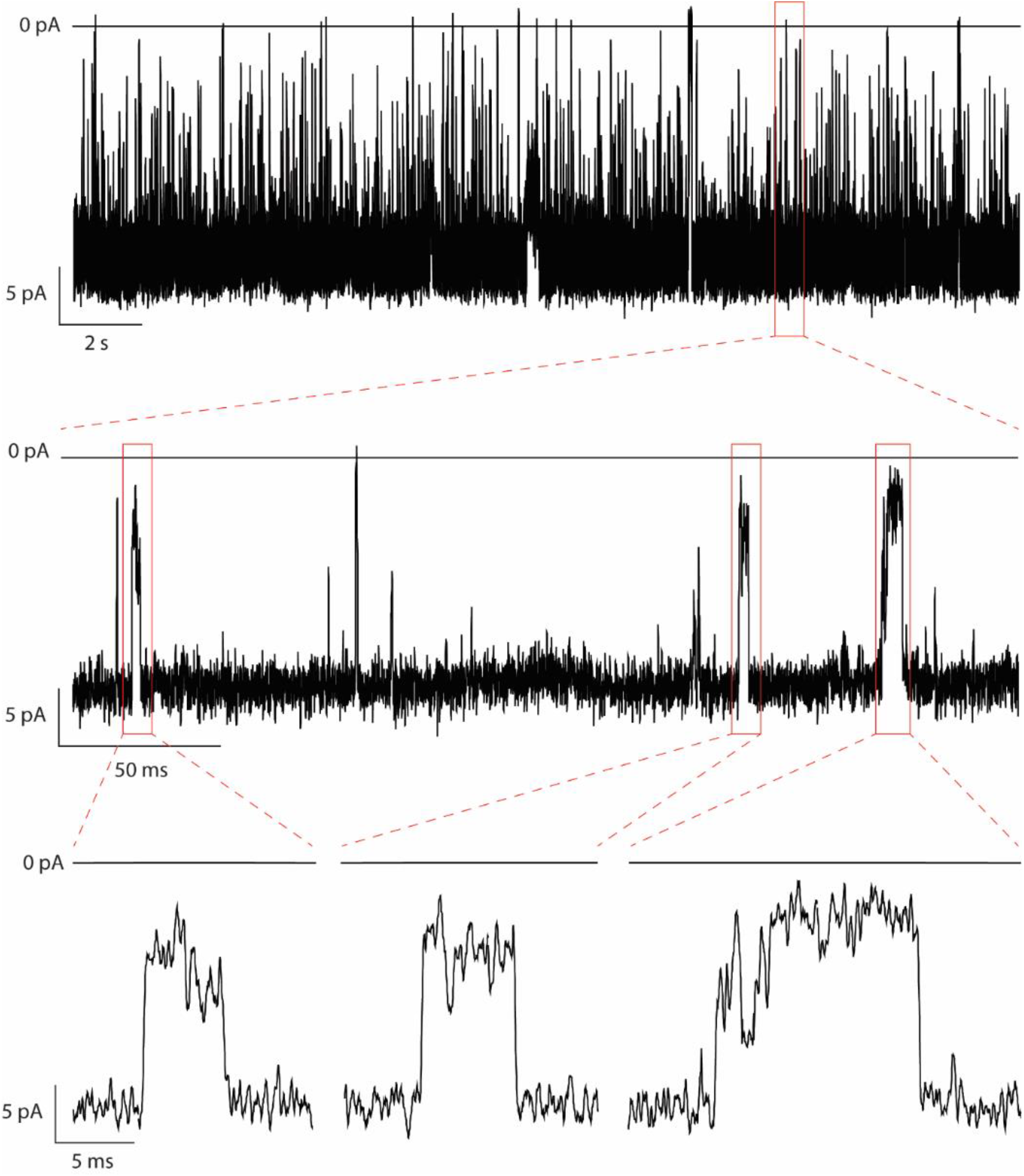
Representative traces of GFP translocation though inactivated proteasome-nanopore at -30 mV and 40°C in 2 mM ATP, 1 M NaCl, 15 mM Tris, pH 7.5. The current signals were filtered at 10 kHz, and sampled at 50 kHz. The traces were then filtered digitally with a Gaussian low-pass filter with a 5 kHz cut-off.

**Supplementary Fig. 19.**
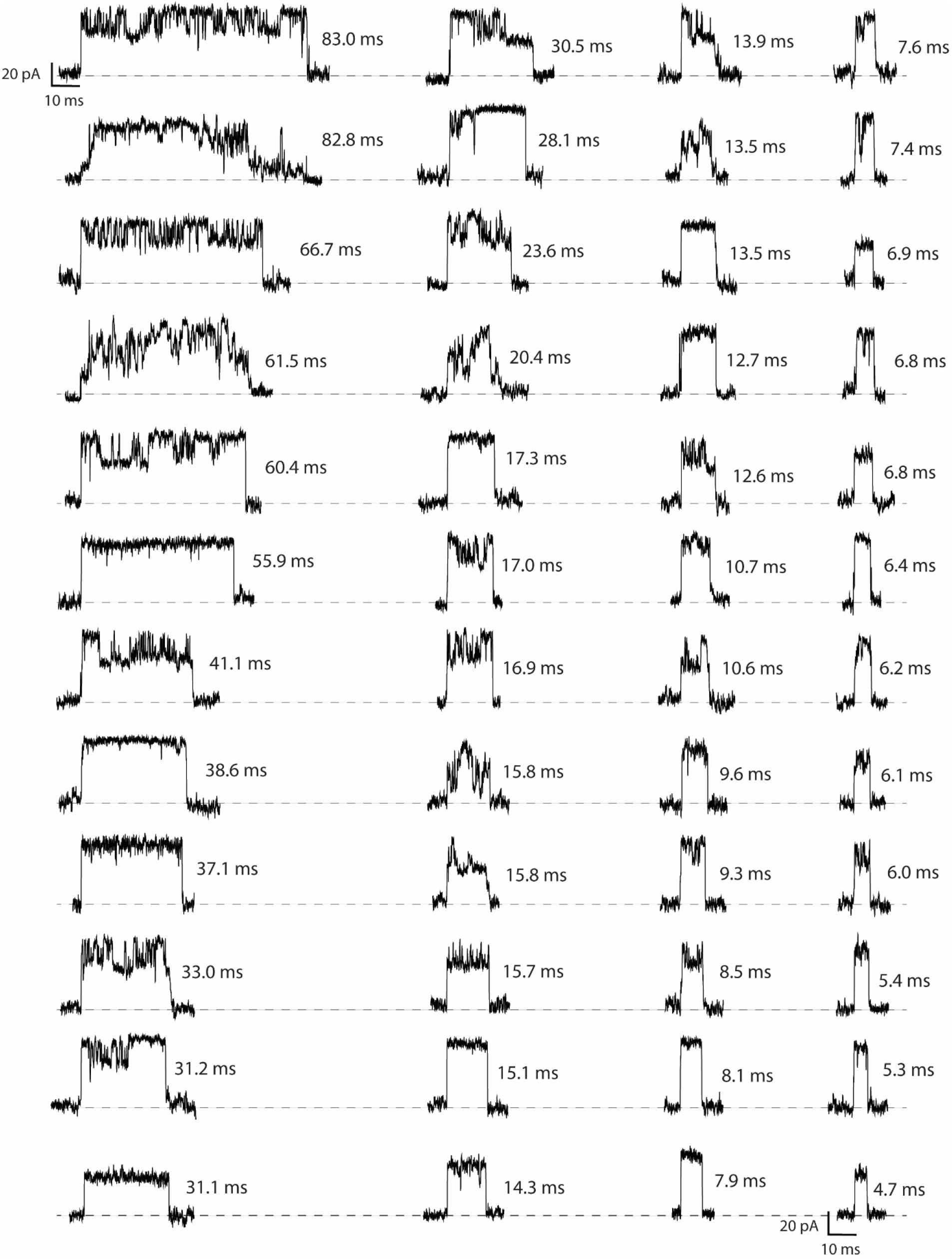
Representative blockades of the GFP translocation though inactivated proteasome-nanopore in the presence of 5 µM VATΔN at 40°C in 2 mM ATP, 1 M NaCl, 15 mM Tris, pH 7.5. The current signals were recorded at -30 mV, filtered at 10 kHz, and sampled at 50 kHz. The traces were then filtered digitally with a Gaussian low-pass filter with a 5 kHz cut-off.

**Supplementary Fig. 20.**
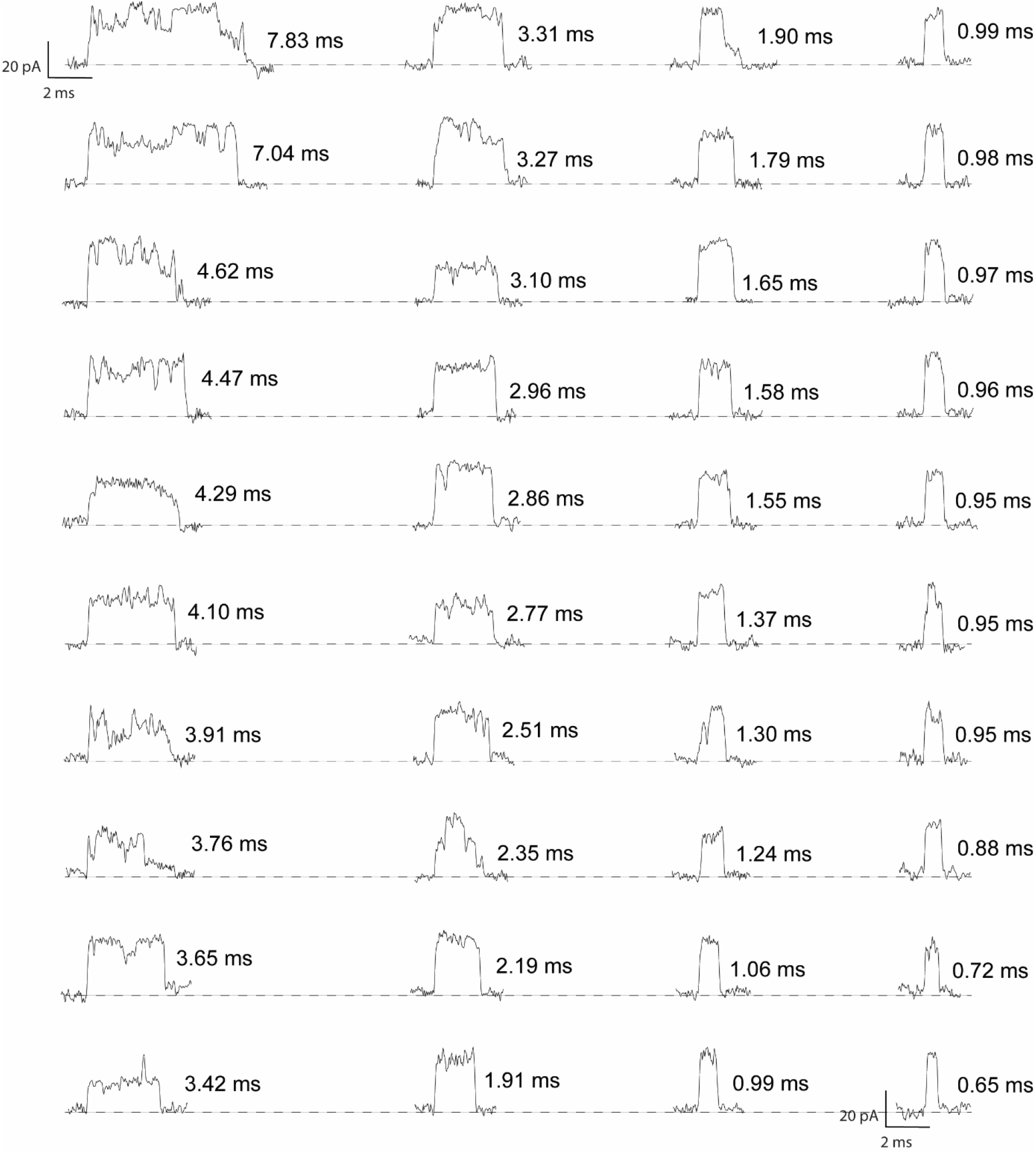
Representative blockades of the GFP translocation though inactivated proteasome-nanopore at 40 °C in 6 mM ATP, 1 M NaCl, 15 mM Tris, pH 7.5. The current signals were recorded at -30 mV, filtered at 10 kHz, and sampled at 50 kHz. The traces were then filtered digitally with a Gaussian low-pass filter with a 5 kHz cut-off.

**Supplementary Fig. 21.**
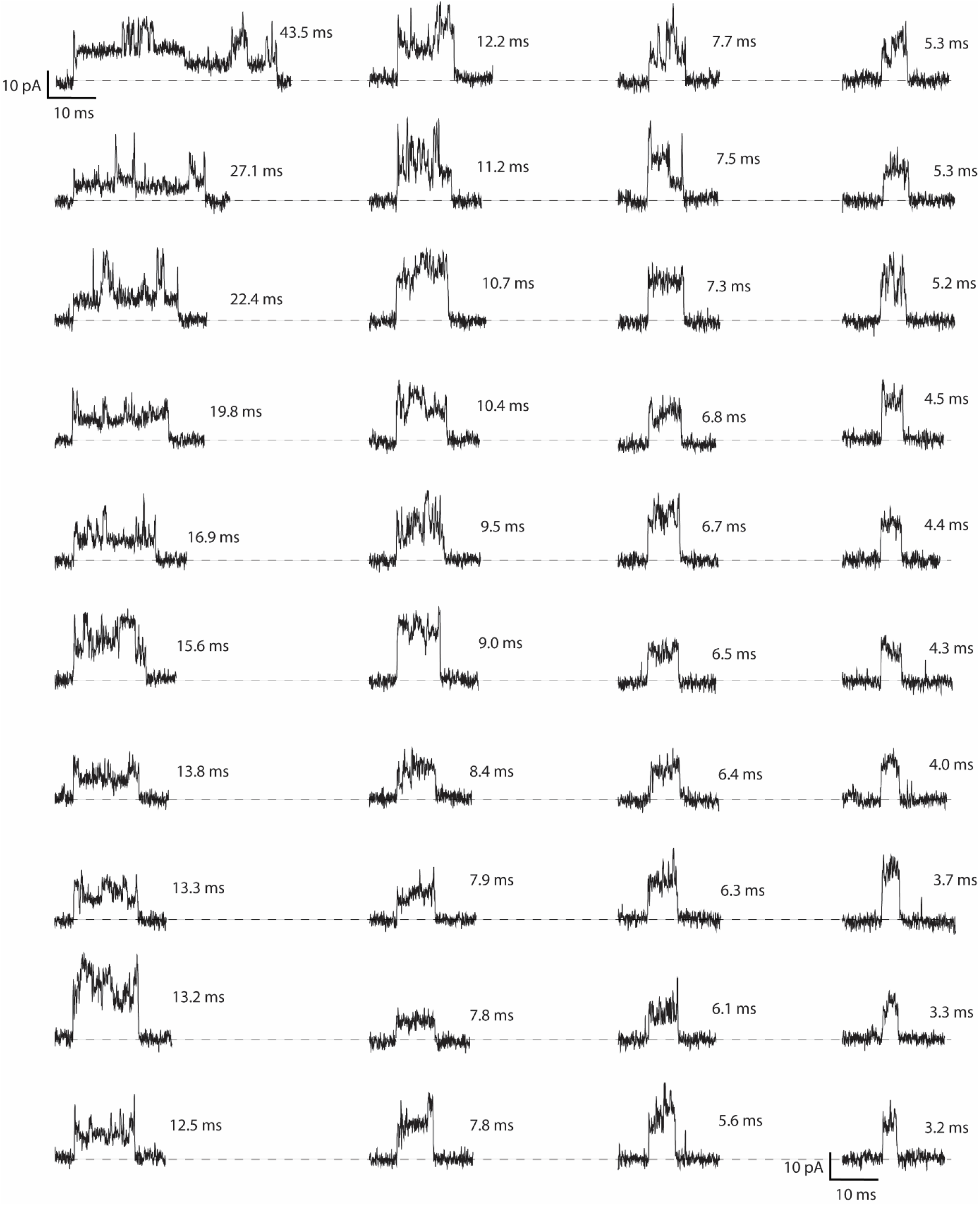
Representative blockades of the GFP translocation though inactivated proteasome-nanopore at 40°C in 1.0 M urea, 2 mM ATP, 1 M NaCl, 15 mM Tris, pH 7.5. The current signals were recorded at -30 mV, filtered at 10 kHz, and sampled at 50 kHz. The traces were then filtered digitally with a Gaussian low-pass filter with a 5 kHz cut-off.

**Supplementary Fig. 22.**
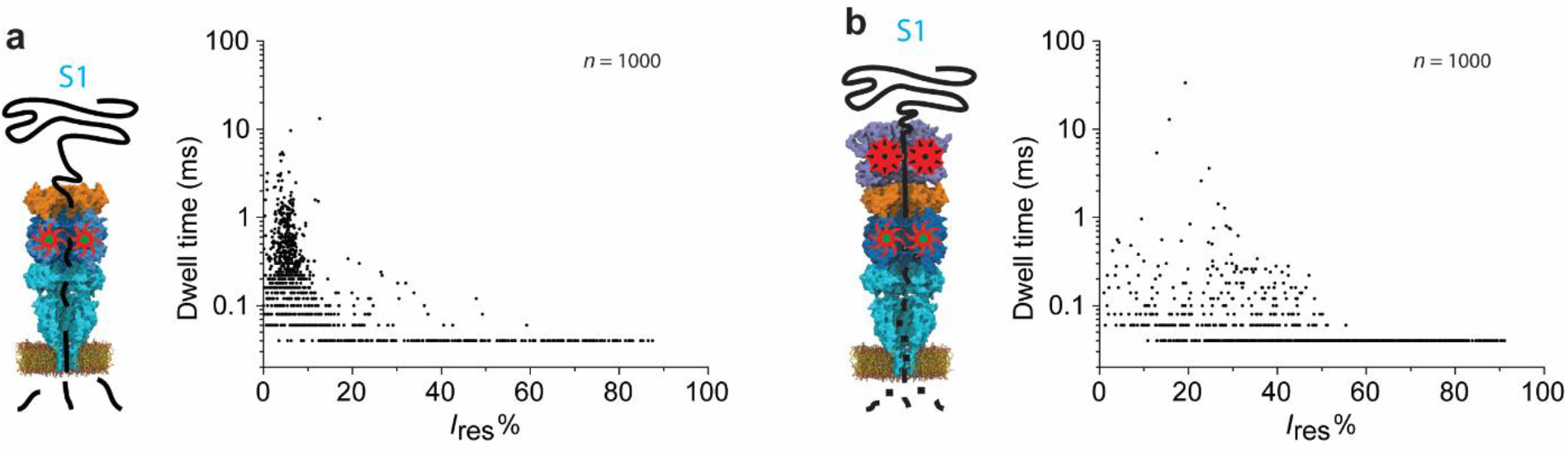
The scatter plots of *I*_res_% versus dwell time provoked by the translocation of substrate 1 (S1, 20 µM) through an active proteasome-nanopore in the absence (a) or presence (b) of VATΔN. Data were collected at 40 °C and -30 mV in 1 M NaCl, 15 mM Tris, pH 7.5, using a 10 kHz low-pass Bessel filter with a 50 kHz sampling rate.

**Supplementary fig. 23.**
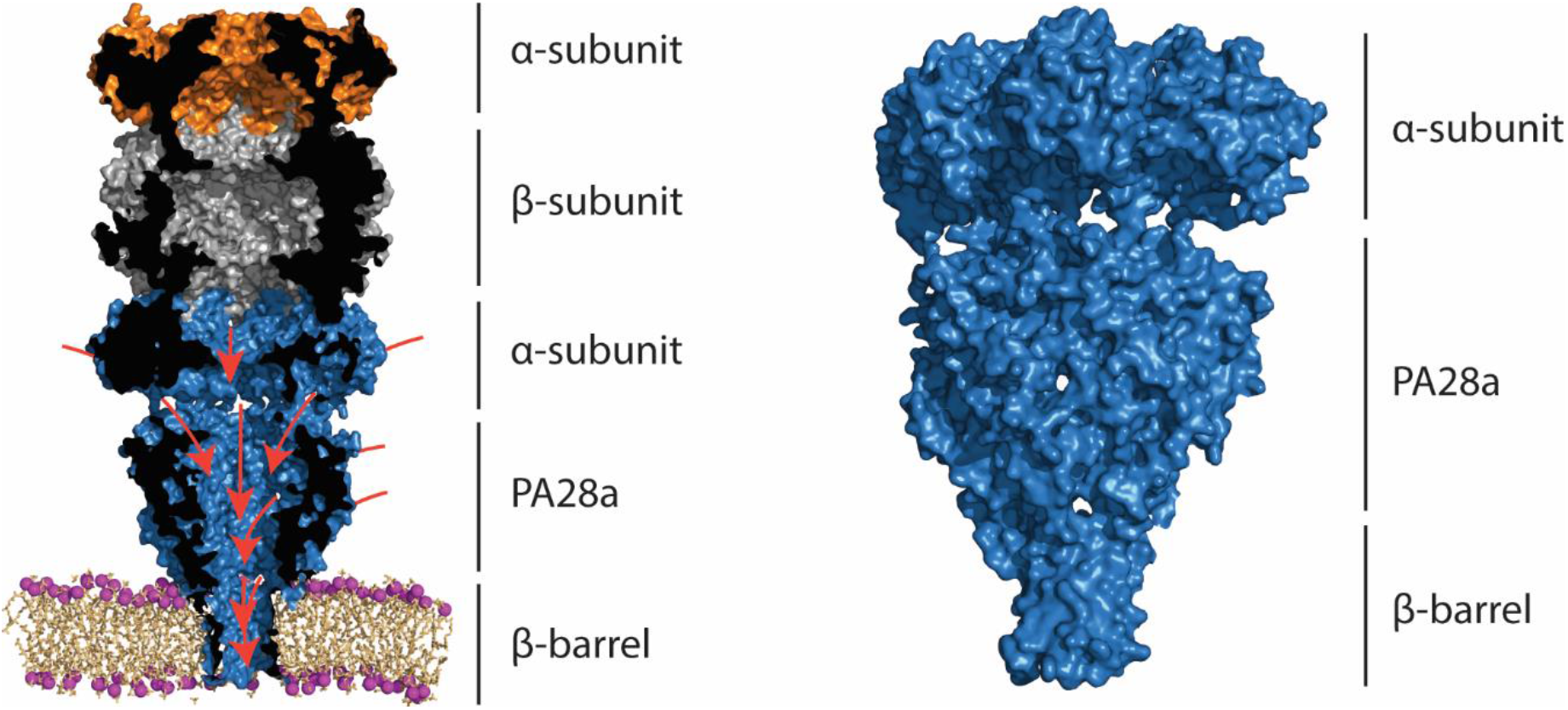
The speculative ion paths. Several potential paths exist that can lead the passage of ions as shown by the proteasome-nanopore structure tested by MD simulations.

